# Low ribosomal RNA genes copy number provoke genomic instability and chromosomal segment duplication events that modify global gene expression and plant-pathogen response

**DOI:** 10.1101/2020.01.24.917823

**Authors:** Ariadna Picart-Picolo, Stefan Grob, Nathalie Picault, Michal Franek, Thierry halter, Tom R. Maier, Christel Llauro, Edouard Jobet, Panpan Zhang, Paramasivan Vijayapalani, Thomas J. Baum, Lionel Navarro, Martina Dvorackova, Marie Mirouze, Frederic Pontvianne

## Abstract

Among the hundreds of ribosomal RNA (rRNA) gene copies organized as tandem repeats in the nucleolus organizer regions (NORs), only a portion is usually actively expressed in the nucleolus and participate in the ribosome biogenesis process. The role of these extra-copies remains elusive, but previous studies suggested their importance in genome stability and global gene expression. Because the nucleolus is also a platform for nuclear organization, we tested the impact of a decreased amount of rRNA gene copies on the *Arabidopsis thaliana* 3D genome organization and stability, using an *A. thaliana* line only containing 20% of rRNA gene copies (20rDNA line). Compared to the wild-type Col-0, the 20rDNA line shows several signs of genomic instability, such as variations in 3D genome organization, spontaneous double-strand breaks accumulation, transcriptomic changes, and higher DNA methylation level. Strikingly, using genomic and microscopic approaches, we identified seven large tandem duplications in direct orientation (TDDOs) ranging from 60 kb to 1.44 Mb. As a consequence, more than 600 genes were duplicated, often associated with an increase in their expression level. Among them, we found several upregulated genes involved in plant-pathogen response, which could explain why the 20rDNA line is hyper-resistant to both bacterial and nematode infections. Finally, we show that the TDDOs create gene fusions and/or truncations and we discuss their potential implications on plant genome evolution.

## INTRODUCTION

In most eukaryotes, hundreds of ribosomal RNA (rRNA) genes compose the nucleolus organizer region (NOR). In *Arabidopsis thaliana Columbia* ecotype (Col-0), 375 tandem 45S rRNA gene copies are located at the top of both chromosomes 2 (NOR2) and 4 (NOR4) (Copenhaver and Pikaard 1996). Only a portion of these copies is actively transcribed in the nucleolus to produce ribosomes. Most of rRNA genes indeed remain transcriptionally inactive and accumulate repressive chromatin modification marks (Grummt and Langst 2013; Pontvianne et al. 2013, 2012, 2010). As in many species, rRNA genes copy numbers are highly variable among *A. thaliana* populations (Kobayashi 2011; Dopman and Hartl 2007; Gibbons et al. 2015; Rabanal et al. 2017). In natural inbred lines found in Sweden, rRNA copy number heterogeneity can account for up to 10 % of genome size variation (Long et al. 2013). Worldwide, *A. thaliana* ecotypes can be found with a rRNA gene copy number ranging from 500 to 2500 in haploid cells (Long et al. 2013). Therefore, 500 copies could be considered as the lowest rRNA gene copy number found *in natura* (Rabanal et al. 2017). In plants, the impact of this variability is not known, but in budding yeast and in *Drosophila*, previous studies suggest that a minimum amount of inactive rRNA genes is necessary for global genome stability. One to two hundred rRNA gene units are usually found in budding yeast, but genome engineering allowed the creation of viable yeast lines with only 40 rRNA gene units. Similarly, shifts in rRNA gene copy number affect genome-wide chromatin marks and alter gene expression in flies (Paredes et al. 2011; Lemos et al. 2008).

FASCIATA (FAS) 1 and 2 are part of the CAF complex required for proper deposition of histones H3 and H4 upon DNA replication (Ramirez-Parra and Gutierrez 2007). In *A. thaliana*, their knock-outs induce a drastic change in rRNA genes copy number, leading to a global reorganization of epigenetic states and the sub-nuclear positioning of the NORs (Pontvianne et al. 2013; Mozgova et al. 2010). Interestingly, crossing *fas1-4* and *fas2-4* mutants and subsequent inbreeding by self-fertilization led to the creation of two *A. thaliana* lines displaying wild-type *FAS* genes with only 20% of the amount of rRNA gene copies in comparison to WT Col-0 (Figure 1A) (Pavlistova et al. 2016). These two 20rDNA lines (hereafter named 20rDNA L6 and L9) have a wild-type phenotype and retained a low amount of rRNA genes for 5 generations (Pavlistova et al. 2016). In this study, we took advantage of this plant material to test the impact of NOR size on 3D genome organization. We also analysed how genome stability and global gene expression was affected in the 20rDNA lines, which revealed unexpected consequences on the genome structure and stability.

**Figure 1:**
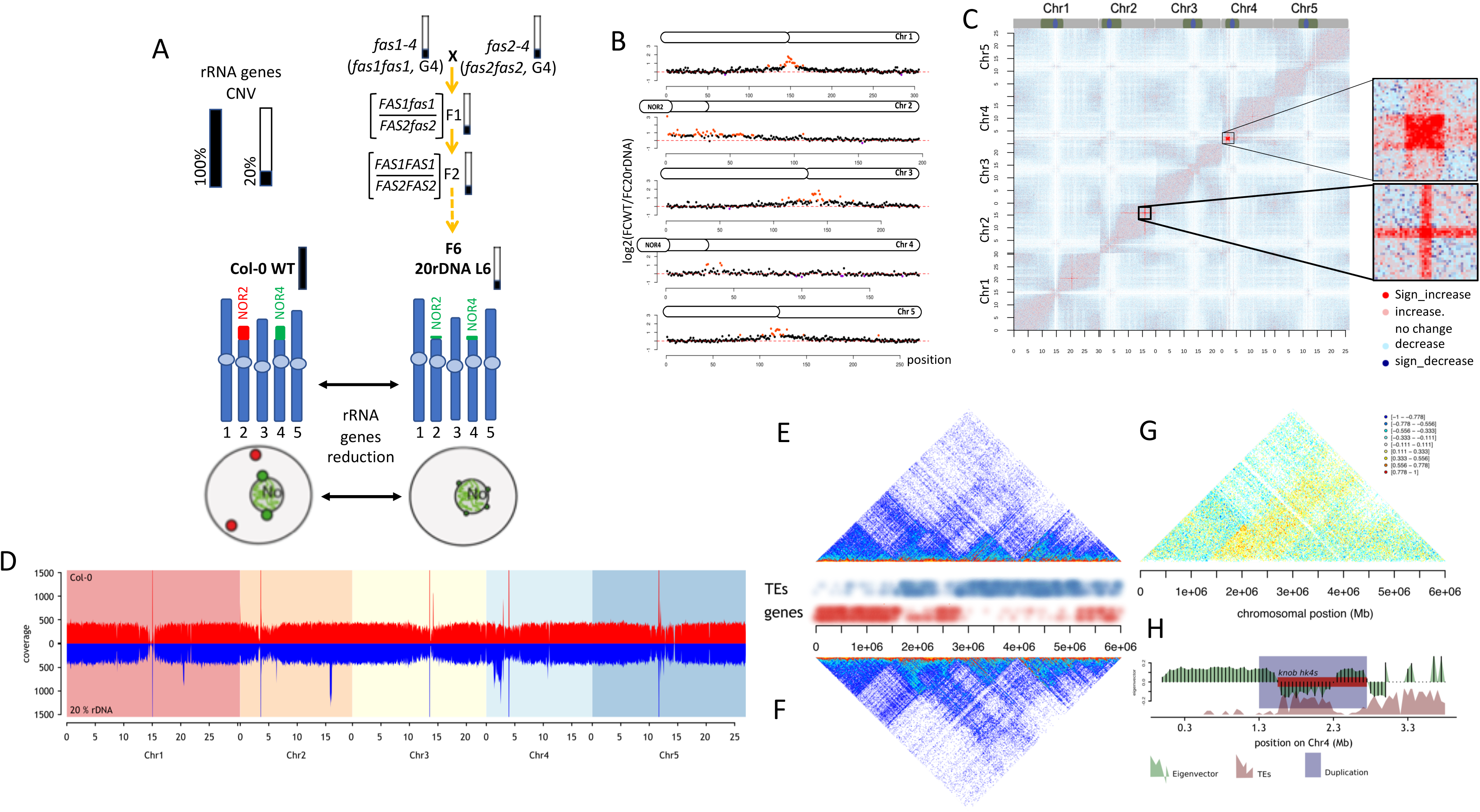
Affected 3D genome organization in the 20rDNA L6F6. **A**. Schematic representation of the obtaining of the 20rDNA L6F6 and its relative content in rRNA gene copies. NOR2 and NOR4 rRNA gene copies are both affected by the reduction and according DNA-FISH experiment, their nuclear distribution in the nucleus changes compare to WT, since all rRNA gene copies associate with the nucleolus (15). **B**. Chromosome plots displaying the relative enrichment of a given genomic segment with the nucleolus. The y-axis displays the fold change nucleolus enrichment between wild-type Col-0 and the 20rDNA line 6. Each dot represents a 100 kb window. Nucleolus-enriched genomic regions above the threshold are colored in red, while depleted regions are colored in violet. **C**. Coverage normalized T-test difference matrix (50 kb bins). The color of each pixel of the matrix is defined by the result of a T-test using the triplicate contact frequencies from wild type and 20rDNA coverage normalized Hi-C samples, respectively. (alternative: T-test difference matrix (50 kb bins). The color of each pixel of the matrix is defined by the result of a T-test using the triplicate contact frequencies from wild type and 20rDNA Hi-C samples, respectively. The two magnified area correspond to the two regions displaying the highest level of contact frequency changes **D.** Summed coverage over triplicate Hi-C data sets in 50 kb bins. The coverage was calculated based on unpaired single end raw Hi-C reads. **E-F** Non-normalized Hi-C snap shot showing the contact frequencies on the short arm of chromosome 4 in WT Col-0 (**E**) *versus* the 20rDNA L6F6 (**F**). TEs and genes are annotated to illustrate the occurrence of euchromatin and heterochromatin, respectively. **G.** Ratio between Hi-C contact frequencies from wild-type and 20rDNA L6F6. Negative ratios: More contacts in the wild-type, Positive ratios: More contacts in the 20rDNA L6F6. **H.** Eigenvector of the Col-0 WT Hi-C data set and annotation of the duplication affecting the knob hk4s. Note the central duplication breakpoint exactly coincides with a change between LSD and a CSD.

## RESULTS

### Low rDNA copies impact NADs composition and 3D genome organization

The nucleolus plays an important role in the spatial organisation of the chromosomes (Bersaglieri and Santoro 2019). Nucleolus-associated chromatin domains (NADs), essentially composed by repressed chromatin domains, localize at its periphery (Nemeth et al. 2010; van Koningsbruggen et al. 2010; Pontvianne et al. 2016b). As rRNA gene nuclear distribution seems to have an important impact in NADs identity both in plant and animal cells (Pontvianne et al. 2016b; Picart-Picolo et al. 2019; Quinodoz et al. 2018), we analyzed NADs composition and 3D organization in an *A. thaliana* line containing only 20 % of rRNA genes named 20rDNA line 6. The 20rDNA line 6 was obtained from the cross between *fas1-4* and *fas2-4* mutant lines as described in (Figure 1A) and is therefore free of T-DNA transgenes (Pavlistova et al. 2016). The 5^th^ generation of the 20rDNA L6 (20rDNA L6F5) was transformed with a transgene expressing ectopically the FIBRILLARIN 2 nucleolar protein fused to the Yellow Fluorescent Protein (FIB2:YFP). Using the FIB2:YFP nucleolar marker, we isolated nuclei and nucleoli from the transformants and identified NADs from the recovered nuclear and nucleolar DNA as previously described (Pontvianne et al. 2016a; Carpentier et al. 2018). Previous studies have clearly demonstrated that in wild-type Col-0 leaf cells, NOR4-derived rRNA genes are expressed and associate with the nucleolus. Conversely, NOR2 is excluded from the nucleolus and NOR2-derived rRNA genes are silent (Chandrasekhara et al. 2016; Pontvianne et al. 2013). As a result, NADs are essentially distributed in the entire short arm of chromosome 4, which juxtapose the active NOR4, associate with the nucleolus (Pontvianne et al. 2016b). Compared to the wild type, NADs in the sixth generation of the 20rDNA L6 line (L6F6) contained both chromosome 2 and 4 short arms (Figure 1B). Among the 434 genes that gained nucleolar association, 144 belong to chromosome 2 (33%). In contrast, chromosome 4 NADs remain stable since only 19 genes gain nucleolus association. These results are consistent with the rDNA transcriptional state, as all left-over NOR2 and NOR4-derived rRNA genes are actively transcribed and associate with the nucleolus (Pavlistova et al. 2016). As in wild type, sub-telomeric regions remain associated with the nucleolus in the 20rDNA L6F6 line (suppl. Figure 1A-B). In total, NADs identification in 20rDNA L6F6 revealed that 5,6 Mb of chromatin domains mainly enriched in silent epigenetic marks changed their subnuclear distribution, which suggests a substantial reorganisation of the nuclear genome. To get a global view of the epigenome 3D organization, we analysed all chromatin-chromatin interactions using chromosome-conformation capture sequencing (Hi-C) (Figure 1C).

We generated triplicate Hi-C samples from both wild-type and 20rDNA L6F6 14-day-old seedlings, allowing us to perform robust statistical analyses (Suppl. Figure 2). We developed a method to assess differences between two given sets of Hi-C samples statistically. To date, comparisons between Hi-C data sets are often unsatisfactory, as they rely merely on visual inspection of relative difference matrices. Making use of our triplicate Hi-C data sets, we performed student T-tests on each contact frequency (pixel of the Hi-C matrix) and determined whether contact frequencies significantly changed between the wild-type and the 20rDNA L6F6 line (Figure 1C and Figure 1E-G). We found several regions displaying a significant (p < 0.01) increase of contact frequencies at several chromosomal locations (Figure 1C, Sign_increase signals). The largest region represents 1.44 Mb, spanning the heterochromatic knob on the short arm of chromosome 4 (*hk4s*), a large heterochromatic region outside the peri-centromeres, and an euchromatic region distal to the knob. Additional regions of increased contact frequencies are present on other chromosomes: 200 kb around the 20 Mb position of chromosome 1; 370 kb around the 15,7 Mb position of chromosome 2; and 60 kb around the 12,5 Mb of chromosome 5. Contact frequencies assayed by Hi-C can be used to detect chromosomal rearrangements (Himmelbach et al. 2018). We suspected that these increased interaction frequencies observed in 20rDNA L6F6 were not primarily due to significant changes in 3D folding principles of the regions, but rather to large duplications events. A duplication would lead to a 2-fold increase in coverage of the affected region and, thus, doubling of interaction frequencies at this region. Indeed, by analysing the genome-wide coverage using unpaired raw Hi-C sequencing reads, we found a significant increase in coverage in all affected regions (Figure 1D). We subsequently normalized our Hi-C matrices for the assayed coverage, which led to the removal of nearly all significantly different contact frequencies stemming from the duplicated regions. However, difference analysis in coverage-normalized Hi-C data showed that short-range contact frequencies within the duplicated regions are significantly depleted. Whether this depletion of contact frequencies is biologically significant or represents an artefact of the normalization procedure is extremely difficult to determine.

To further examine potential differences in 3D folding principles between wild-type Col-0 and 20rDNA L6F6, we performed a principal component analysis (PCA) to retrieve the Eigenvector, which is characteristic of 3D-folding patterns of a Hi-C data set (Grob et al. 2014; Lieberman-Aiden et al. 2009). Sign-changes in the Eigenvector delineate basic 3D folding domains, known as Loose Structural Domains (LSDs) and Closed Structural Domains (CSDs), which are analogous to animal A and B compartments (Lieberman-Aiden et al. 2009). We could not identify significant changes in the Eigenvectors between the two lines that would go beyond the variation observed within the lines. We then calculated the contact density for each genomic bin, which is determined by the number of contact frequencies observed within a given region surrounding the genomic bin of interest. In line with our difference analysis, we could observe a decrease in contact densities in duplicated regions, however, no significant difference were observed elsewhere in the genome.

Subsequently, we were analyzed whether the location of the duplications might have been biased based on *a priori* folding principles in the wild type. Hence, we compared duplication breakage points with the Eigenvector obtained by the PCA analysis of the wild-type Hi-C data (Suppl. Figure 3). Intriguingly, we observed that in a majority of duplications at least one of the breakage points coincides with sign-changes (CSDs to LSDs) or directional changes (valleys and peaks within a structural domain) in the Eigenvector. Hence, the changes in 3D-conformation may have facilitated the occurrence of duplications. For the knob *hk4s*, this was most prominent, where the more central breakpoint exactly co-localizes with the change between the CSD and the LSD, which defines the knob border in 3D. This duplication exactly co-localizes with the ancient inversion breakpoint that gave rise to the knob (Figure 1H) (Zapata et al. 2016). This suggests the existence of continuously fragile chromosomal regions, the borders between structural domains being diagnostic for these regions.

### Characterization and impact of duplication events on gene expression

To confirm and precisely map the duplicated regions in the 20rDNA L6F6 line, we performed long-read resequencing using Nanopore technology. We obtained 6,4 Gb of total sequences with a mid-size of 6 kb. We then analyzed the sequencing coverage against the TAIR10 *A. thaliana* Col-0 reference genome to identify the highly covered regions (Figure 2A). We confirmed the existence of the four large duplications found by Hi-C. We also identified three additional minor duplications: a 80 kb-long duplication around the 6,4 Mb position of chromosome 2, a 60 kb-long duplication around the 10,7 Mb position on chromosome 4, and an additional duplication of 100 kb in the already duplicated region on the *hk4s* (Figure 2A). These seven duplication events correspond to tandem duplications in direct orientation (TDDO) and were named TDDO1 to TDDO7. Combined, they correspond to a gain of 2,31 Mb per haploid genome. These TDDO provoke copy number variations (CNV) of 626 genes and 851 Transposable Elements (TEs) in the 20rDNA L6F6 line. In parallel, we also confirmed a significant reduction of reads mapped to rRNA genes in the 20rDNA L6F6 line.

**Figure 2:**
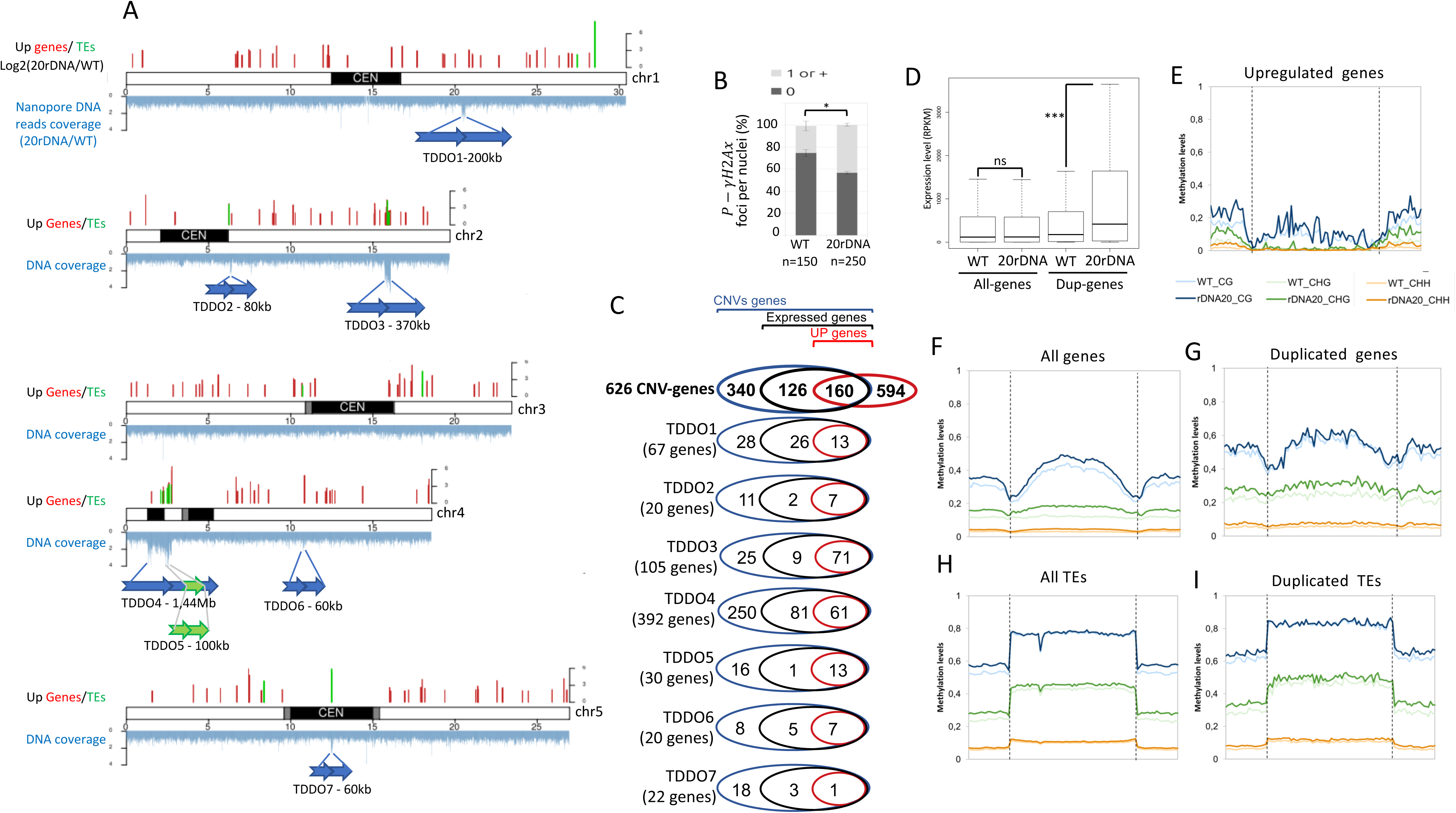
Identification of large tandem duplication in direct orientation and characterization of their transcriptome and epigenome. **A.** Distribution of the reads obtained by nanopore sequencing along the chromosomes in the 20rDNA line 6 compare to wilt-type Col-0 (bottom_blue). Upregulated genes (red bars) and TEs (green bars) with an adjusted *p* value < 0.01 and a log2FoldChange > 2 (top). The blue arrow represents the localization and the orientation of the TDDO identified in the 20rDNA line 6. **B.** Histogram showing the percentage of nuclei displaying at least one phosphorylated γH2Ax foci in WT Col-0 *versus* 20rDNA line 6. **C.** Venn diagrams representing the proportion of expressed genes (containing at least 2 reads/genes in wild-type) and upregulated genes (*p* value < 0.01, FC>2) among all the duplicated genes or in each TDDOs. **D**. Dot-plot revealing the relative expression of all genes or duplicated (DUP) genes in 3 week-old leaves in WT Col-0 or in the 20rDNA line 6. *p* value = 0,0005 (***) calculated using a Wilcoxon test. (**E-I).** global DNA methylation analyses from genome-wide bisulfite sequencing experiments in WT col-0 versus the 20rDNA line 6. Global CG, CHG and CHH methylation is shown for upregulated genes with an adjusted *p* value < 0.01 and a log2FoldChange > 1.5 (E), all genes (F), duplicated genes (G), all TEs (H) and duplicated TEs (I).

The occurrence of duplication events is a sign of genomic instability. Thus, the chromosomal rearrangements observed in the 20rDNA L6F6 could be the consequence of double-stranded breaks (DSBs). To test this hypothesis, we compared the amount of spontaneous DSBs between 20rDNA L6F6 and wild-type Col-0 cell by performing immunostaining of serine 139-phosphorylated H2AX histone variants (P-γ-H2Ax), which is a marker of (Charbonnel et al. 2010). P-γ-H2Ax foci were detected at a higher rate in 20rDNA L6F6 nuclei compared to wild-type nuclei (Figure 2B and Suppl. Figure 4A). Accumulation of DSB foci can potentially be associated with a DNA repair defect. This hypothesis is supported by an increased susceptibility of the 20rDNA L6F6 line to a treatment with the genotoxin bleomycin (Suppl. Figure 4B).

Changes in the organization of the 3D genome as well as CNVs can have an impact on chromatin marks and gene expression. We therefore analysed the global gene expression pattern by polyA+ RNA-seq and the methylome by whole-genome bisulfite sequencing (WGBS) in wild-type Col-0 *versus* 20rDNA L6F6 line. Four replicates per samples of RNA-seq analyses identified differentially accumulating transcripts: 321 upregulated genes and 14 upregulated TEs, as well as 37 downregulated genes but no downregulated TE in the 20rDNA L6F6 line (with an adjusted p-value <0.01 and log2(FoldChange) >1.5 or <1.5) (Figure 2A and Suppl. Figure 5A). We confirmed these results using quantitative RT-PCR (RT-qPCR) on 9 randomly chosen genes and TEs (Suppl. Figure 5B). We did not find any correlation between differentially expressed genes and genes located in the NAD of 20rDNA L6F6 line (Figure 2A; Suppl. Figure 6A-C).

However, we found that duplicated genes and TEs were significantly more expressed (Suppl. Figure 6D-E). 57% (8) of the upregulated TEs are also duplicated (Suppl. Figure 6E). If we consider the 321 upregulated genes with a foldchange enrichment of 1.5, we found that 22% of these genes (71) belonged to duplicated genes, while the TDDOs only represent 2% of the genome. Conversely, no overlap was found between duplicated and down-regulated genes. Higher expression can only be observed from initially expressed genes in wild-type plants. Only 286 duplicated genes are actually expressed and 160 of them are at least twice more expressed than in wild-type (Figure 2C). Interestingly, depending on their genomic location, TDDOs perform differently. For instance, most of the TDDO3 genes accumulates at least twice more transcripts in the 20rDNA L6F6 (71 upregulated genes out of the 80 expressed genes), while genes present in the TDDO4 that contain the heterochromatic *hk4s* region are less upregulated (61 foldchange>2 genes out of the 142 expressed genes) (Figure 2C). Finally, box-plot analyses of all genes *versus* the duplicated genes indeed revealed their overall ability to over-accumulate more transcripts in the 20rDNA L6F6 line (Figure 2D). Thus, our data strongly suggest that duplication leads to an increase expression.

To analyse the impact of CNVs at the DNA methylation level, we performed whole-genome bisulfite sequencing (WGBS) in wild-type Col-0 *versus* 20rDNA L6F6 line in triplicate. We observed a modest increase in CG methylation in the 20rDNA L6F6 line at genes but not at TEs (Figure 2F, 2H). This increase is also modest when considering duplicated genes only (Fig 2G). We also found an increase in CG, CHG and CHH methylation in the 5’ and 3’ sequences flanking the genes (Figure 2E-G; Suppl. Figure 7). At TEs, methylation was affected in both CHG and CHH contexts, but not in the CG context (Figure 2H-I; Suppl. Figure 8). Although significant changes can be observed at the DNA methylation level, they seem to concern all genes and TEs, and not a specific group of genes. In other words, both duplicated and upregulated genes are concerned by the global DNA methylation increase. Intriguingly, gene body methylation of the upregulated genes was not concerned by this DNA methylation increase (Figure 2E and Suppl. Figure 7). Finally, differentially methylated regions (DMRs) identified in the 20rDNA L6F6 compare to WT Col-0 did not demonstrate a potential overlap between upregulated genes and hypomethylated regions (Suppl. Figure 6A-C).

### Duplication events are linked to a higher pathogen resistance

Gene categories implicated in biotic and stress responses are particularly enriched in the upregulated genes in the 20rDNA L6F6 line (Figure 3A). We performed RT-qPCR analyses and confirmed the overexpression of key genes involved in the plant-pathogen response (Figure 3B). Among these genes are *Pathogenesis-Related* (*PR*) 1 and 5 genes, whose higher expression levels are usually correlated with increased resistance against bacteria and nematodes (Wubben et al. 2008). Notably, some of these genes were found in the duplicated regions of chromosome 2 and 4 (TDDO3 and TDDO4). It is therefore possible that their higher expression rate is a consequence of the duplication events (Suppl. Figure 9). *ASYMETRIC LEAVES 1* (*AS1*) has an evolutionarily conserved role in plant-pathogen interactions (Yang et al. 2008). AS1 indeed acts as a positive regulator of extracellular defenses against bacterial pathogens in a salicylic acid-independent manner (Nurmberg et al. 2007). In addition, genes encoding four Cysteine-rich Receptor-like Kinases (CRKs), located in the duplicated part of the *hk4s*, also overaccumulate transcripts in the 20rDNA L6F6 line (Suppl. Figure 9). Among these genes is *CRK36,* whose overexpression was shown to be sufficient to enhance pattern-triggered immunity response and bacterial pathogen resistance (Yeh et al. 2015).

**Figure 3:**
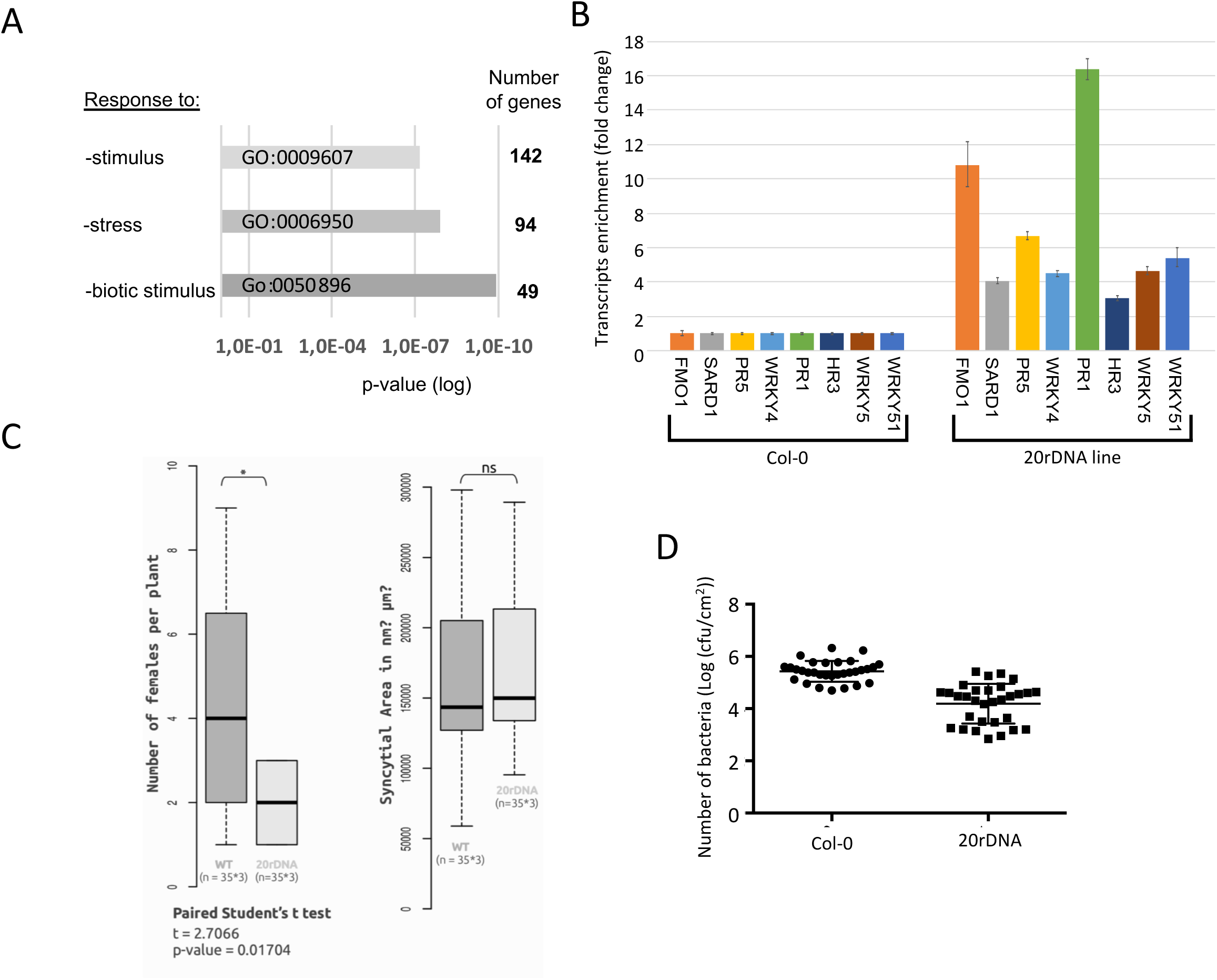
The 20rDNA line 6 overexpressed biotic stress genes and is more resistant to nematod and bacterial pathogenes. **A**. Go term enriched in the upregulated genes pool identified in the 20rDNA line 6. **B**. Histogram displaying the relative transcript enrichment for 8 genes implicated in the biotic stress response using qRT-PCR in WT Col-0 versus the 20rDNA line 6. **C**. WT Col-0 and the 20rDNA line 6 were inoculated with the sugar beet cyst nematode (*Heterodera schachtii*). 4 weeks after inoculation, the number of adult females per plant was determined. Data are the average number of adult females ± SE (n = 35 X3). Data from the 3 independent experiments were pooled and are shown (Left histogram). The relative size of the syncytium cells was measured between both lines but no significant changes were noticed (right histogram). **D**. WT Col-0 and the 20rDNA line 6 were inoculated with Pseudomonas strain DC3000 at 5 x 107 cfu/ml. Relative bacterial growth was determined 3 days after infection and is shown on the plot.

We therefore challenged the 20rDNA L6F6 line and wild-type plants with phytopathogenic agents to test their resistance capabilities. *A. thaliana* is susceptible to various pathogens, from prokaryotes to multicellular organisms. We first tested the ability of the nematode *Heterodera schachtii* to infect both the wild-type Col-0 and the 20rDNA L6F6 line (Figure 3C). We observed that only half the number of females was able to develop on the 20rDNA L6F6 plants in comparison with wild-type Col-0 plants (Figure 3C). However, we did not observe a change in the syncytium size, i.e., the plant feeding structure induced by these nematodes. Secondly, we tested the ability of the 20rDNA L6F6 to be infected by the virulent bacteria *Pseudomonas syringae* strain DC3000. Three days after inoculation, bacterial growth was significantly lower in the 20rDNA L6F6 line (Figure 3D) than in the wild type. In conclusion, we were able to correlate that the higher accumulation of transcripts from genes implicated in the plant-pathogen response is able to increase its ability to resist against at least two types of distinct pathogens.

### Appearance of the duplication events in the 20rDNA line

One of the key questions is when these TDDOs appeared. The 20rDNA L6F6 line was obtained from the cross between *fas1-4* and *fas2-4* mutants that both display low amounts of rDNA copies (Pontvianne et al. 2013; Mozgova et al. 2010). We therefore tested by quantitative PCR (qPCR) the presence of the duplicated regions in the parental lines and in the offspring. We focused on TDDO3 and TDDO4, the two largest genomic duplications (Figure 2A). Our analyses revealed that TDDO4 was already present in the *fas2-4* parent, but not in the *fas1-4*. TDDO4 segregated in the F2 after the cross, being fixed in 20rDNA L6, but was lost in 20rDNA L9 (Figure 4A). Its absence/presence was also confirmed by DNA-Fluorescent *In situ* Hybridization (FISH) (Figure 4B). We used two probes generated from BAC clones: one recognizing a portion of the TDDO4 (*hk4s* – T5H22) and one recognizing an unduplicated genomic region located between the TDDO4 and the NOR4 (F5I10). Different cell-types were analysed from vegetative and reproductive tissues (Figure 4B and Suppl. Figure 10). While a similar number of signals were detected in wild-type Col-0 nuclei for both probes, more signals corresponding to the TDDO4 were detected in the 20rDNA L6F6 nuclei (Suppl. Figure 10). Analyses of pachytene chromosomes clearly demonstrated that the additional signal actually belonged to the same chromosome, which confirmed the duplication hypothesis (Figure 4B). In parallel, the analyses of nuclei from the 20rDNA L9 line confirmed the absence of the TDDO4 at the 5^th^ generation (F5) (Figure 4B and Suppl. Figure 10).

**Figure 4:**
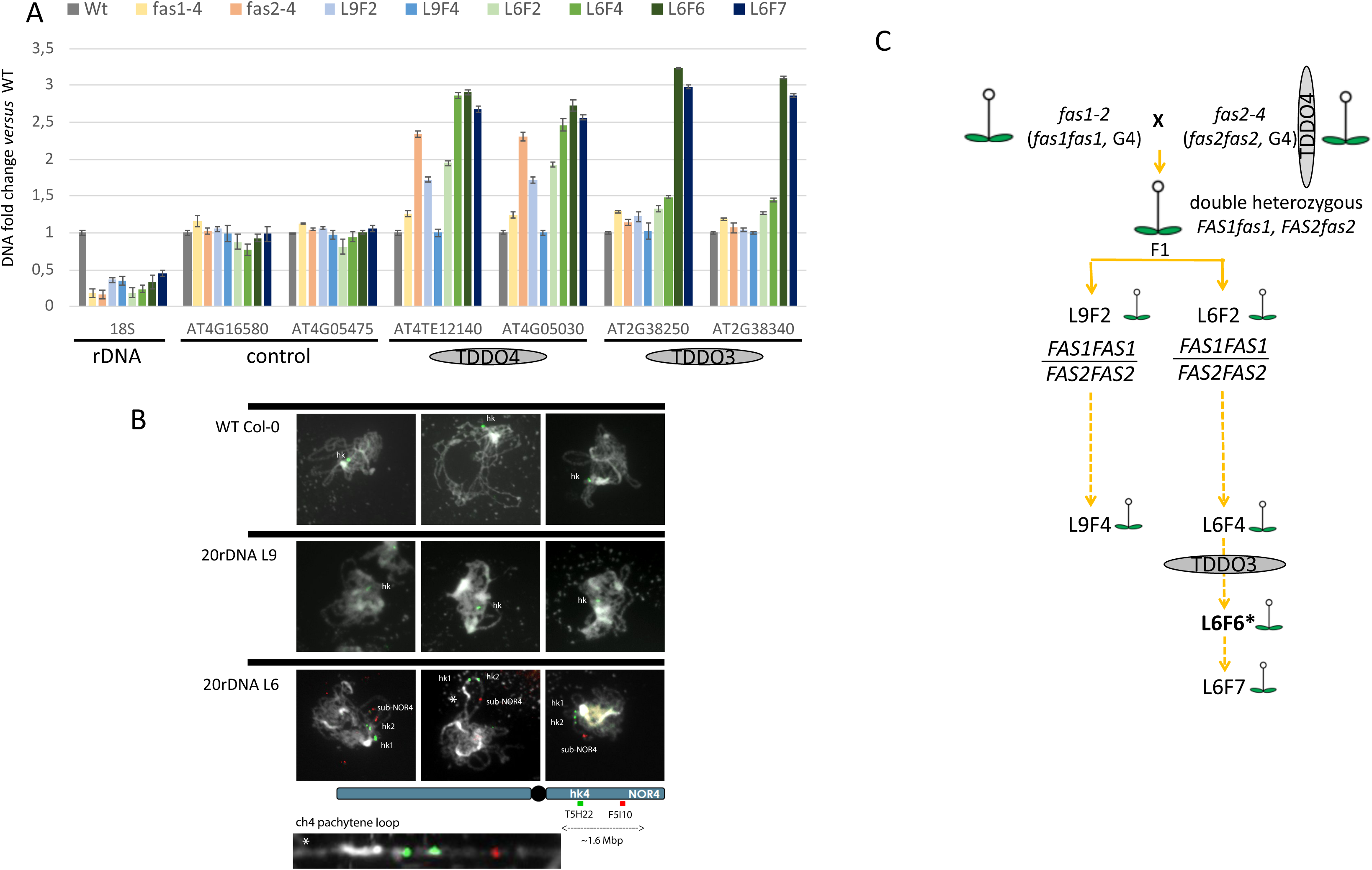
Apparition of the 2 major TDDO events. **A**. CNV of genes present or not in TDDO and of rRNA genes were determine by qPCR. Their relative enrichment was determined in WT Col-0, the parent lines *fas1-4* and *fas2-4* and in 2 of the siblings line 6 and 9 during several generations. Relation between the lines are shown in (C). rRNA genes CNVs was followed *via* probes amplifying the 18S, the *loci* AT4G05475 and AT4G16580 that are not duplicated are controls, while the *loci* AT4TE12140 / AT4G05030 and AT2G38250 / AT2G38340 allow the identification of the 2 largest duplication TDDO4 and TDDO2 respectively. **B.** DNA-FISH analyses of 2 *loci* present on kr4s, distant by 1.6 Mb: one present on the TDDO4 (BAC T5H22 – green) and one absent of the duplicated region (BAC F5I10 – red). Three Pachytene chromosomes from WT Col-0, 20rDNA Line 6 and 20rDNA Line 9 are shown. A zoom of the unlooped kr4s from the middle panel of the 20rDNA line6 is presented at the bottom of the panel. **C.** Schematic representation of the obtaining of the 20rDNA line 6 and 9 and its relative. The putative appearance of the TDDO3 and TDDO4 events are shown.

In addition, qPCR analyses revealed that the TDDO3 is absent from the *fas1-4* and *fas2-4* mutant genomes, and from the first generated offspring. Our data suggest an appearance of the TDDO3 between the 3^rd^ and 6^th^ generation of line 6 (Figure 4A). Late appearance of the TDDO3 is also confirmed by its absence from the 20rDNA L9 line. Together, these analyses suggest that the two TDDOs appeared independently, either in the parental line or in the imbred line resulting from the *fas1-4* and *fas2-4* cross (Figure 4C). Furthermore, the correlation between low rDNA abundance and TDDO appearance remains valid, since *fas2-4* also contains only 20% of the rDNA copies compared to the wild type.

### Duplication events create chimeric genes

Most of the times, TDDOs keep genes intact and do not lead to gene loss. However, truncated genes can be generated at the breakpoint junction, while keeping intact genes on the edges of duplication (Newman et al. 2015). Besides, when breakpoints are located in two different genes in the same orientation, gene fusions can be created if the reading frame is preserved. In the 20rDNA L6F6, we systematically analyzed the TDDO breakpoint junctions (Suppl. Figure 11). Out of the seven cases of TDDO identified in our study, three potentially created fused or truncated proteins (Figure 5A-F). On chromosome 1, the TDDO1 fused the first exon of gene *AT1G55325* that encodes the N-terminal domain of the MEDIATOR 13-like with 4 of the 5 exons of *AT1G54770* that encodes the FCF2 pre-rRNA processing factor. On chromosome 2, although genes are in the opposite orientation, the TDDO3 creates a shorter ORF of the *AT2G38460* gene that potentially produces a truncated FERROPORTIN 1 protein. Finally, on chromosome 4, the TDDO4 fused the *AT4G05475* gene to a TE (*AT4G02960*), leading to the potential expression of three new ORFs, including one that encodes a protein with two Leucin Rich Repeats (LRR) (Figure 5A-F). We then systematically analysed the presence of these chimeric genes in the parental *fas* mutant lines and in the 20rDNA L6 and L9 lines (Figure 5G). The TDDO1-derived chimeric gene was detected in both line 6 and 9, suggesting the appearance of the TDDO1 after the cross between *fas1* and *fas2* (Figure 5G and Suppl. Figure 12). Chimeric genes generated from TDDO3 were specifically detected in line 6 while chimeric genes generated from TDDO4 were detected in *fas2*, line 6 and line 9 plants, confirming the results obtained in Figure 4.

**Figure 5:**
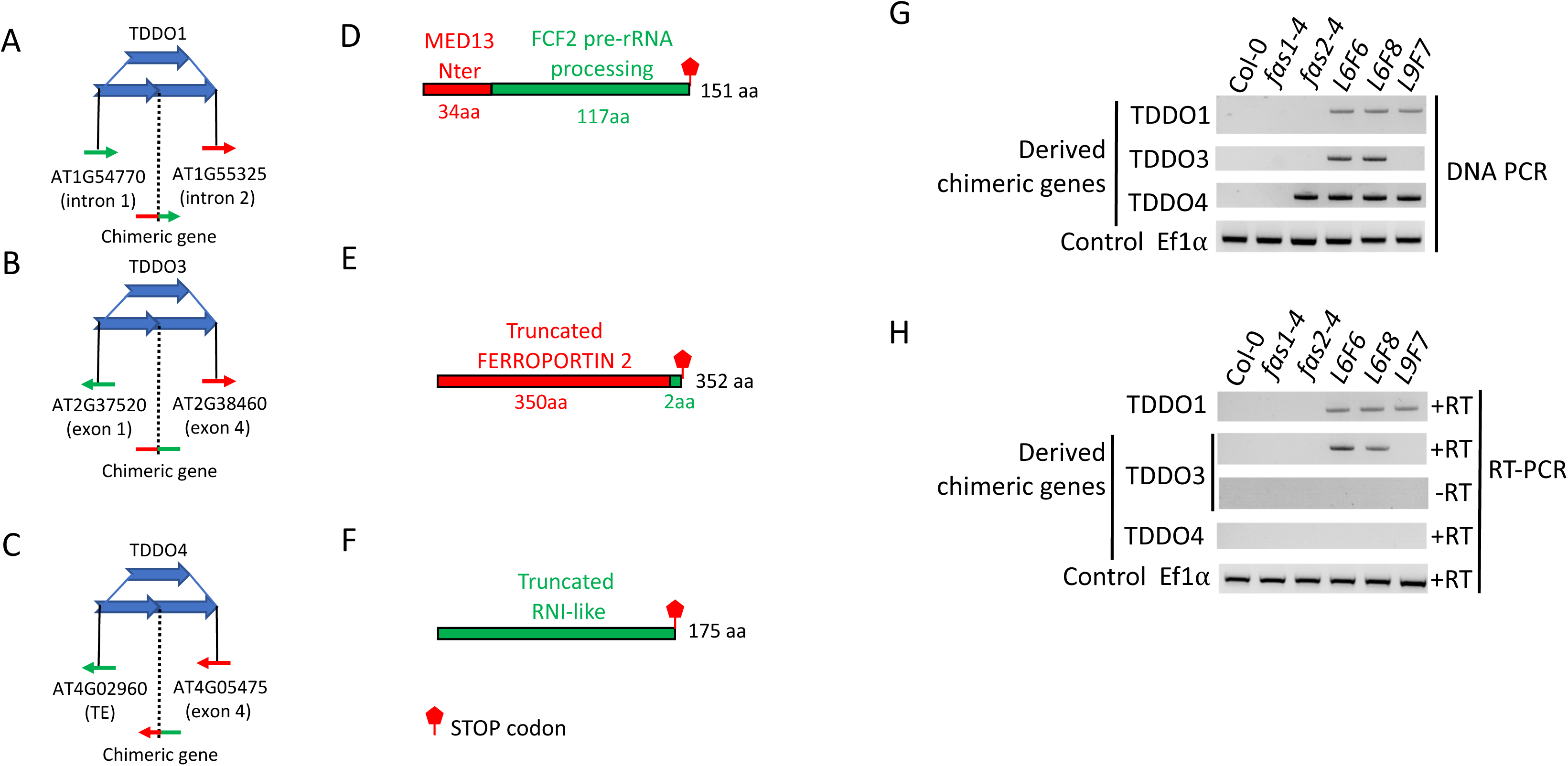
TDDO provoke chimeric genes formation. **A-C**. Schematic representation of the TDDO 1, 3 and 4 that provoked TE and/or gene fusion in the 20rDNA line 6. Genes or TEs present in the breaking junction (and their orientation are shown. Potential fused protein sequences are displayed with the putative nature and their domains. **D-F.** Open Reading Frames (ORFs) potentially generated at the breaking points. The TDDO1 provoke the fusion of the first exon of AT1G55325 that encode for an ATPase motif of MEDIATOR 13 and the last four exons of AT1G54770 that contains an RNA processing domain (D). The chimeric gene created between AT2G37520 and AT2G38460 potentially encode for a truncated FERROPORTIN protein (E). The breaking points at the TDDO4 fuse the 5’ sequence of a TE (AT4G02960) with the second and last exon of the gene AT4G05475 whom sequence encode two Leucine Rich Repeats of AT4G05475 (LRR) (F). **G-H.** PCR performed with primers flanking the breaking junctions of the TDDO1, TDDO3 and TDDO4 in the wild-type Col-0, the two mutants *fas1-4* and *fas2-4*, the 20rDNA lines L6 (generations F6 and F8) and L9 (generation F7). All PCR products sequences were confirmed by Sanger sequencing. Genomic DNA (G) and cDNA (H) were used as template. The locus encoding the elongation factor *EF1α* was used as a loading control.

We finally investigated whether these chimeric genes were transcribed. A first analyses of our RNA-seq data revealed that these genes were all able to accumulate transcripts. Using RT-qPCR, we confirmed the expression of the TDDO1 and 3 derived chimeric genes, as well as the ability of the TDDO1-derived chimeric gene to be properly spliced, gene fusion occurring in the middle of an intron (Suppl. Figure 12). However, although reads could be detected in the RNA-seq data, we did not detect any signals for the TDDO4-derived chimeric gene by RT-PCR (Figure 5H). In conclusion, our data demonstrate that TDDOs can create expressed chimeric genes.

## DISCUSSION

Genomic structural variations shape animal and plant genomes (Krasileva 2019). Within a period of several millions of years, numerous rearrangements have occurred to shape the *Arabidopsis thaliana* genome, including duplications, translocations, inversions and deletions (Blanc et al. 2000; Henry et al. 2006). Recently, genome analysis of seven accessions of *A. thaliana* revealed that they contain on average, 15 MB of rearranged sequences, generating CNVs for thousands of genes (Jiao and Schneeberger 2019). In this case, deletions, gain or loss of copies are considered important sources of CNVs and have potentially occurred in tens of thousands of years of evolution (Fulgione and Hancock 2018). CNVs occurring in the context of tandem duplication events represent between 3 and 4 Mb of genomic sequences in each of the seven accessions sequenced (Jiao and Schneeberger 2019). In our case, only a few generations were necessary to gain 2.3 Mb of genomic sequences by tandem duplications.

The rapid occurrence of these rearrangements is particularly intriguing. The relative sensitivity to genotoxic stress and the detection of a higher rate of spontaneous DSB in our 20rDNA line is certainly one source of their appearance (Figure 2B and Suppl. Figure 4), but the precise mechanisms remain to be determined. One possibility is the implication of non-allelic homologous recombination (NAHR), usually responsible for TDDO (Krasileva 2019; Zhang et al. 2013). This mechanism can generate segmental duplications or deletions. In the 20rDNA L6F6, we detected duplications but no deletion, probably because of their deleterious effects. Two other particular aspects of the detected TDDOs are their large sizes and locations, ranging from 60 kb to 1.44 Mb on four of the five chromosomes. (Figure 2A). The seven TDDO sites do not share any genetic feature, and breakpoint junctions are not enriched in repetitive elements or particular genes (Suppl. Figure 3). However, our Hi-C data revealed that sign-changes in the Eigenvector seem to be overrepresented at breaking junctions, suggesting a potential link between the 3D genome folding and the occurrence of TDDOs. The systematic identification and characterization of additional TDDOs would be necessary to strengthen this hypothesis.

It is also intriguing that five of the seven TDDOs, which represent 89% of the total segmental duplications, are located on NOR-bearing chromosomes (Figure 2A). Due to their tandemly repeated nature, NORs are indeed subjected to an inherent instability (Nelson et al. 2019). Therefore, the existence of a sensing system monitoring their abundance has been proposed (42), potentially *via* unequal sister chromatid exchange (Tartof 1974a, 1974b). The 20rDNA lines derive from the cross between *fas1* and *fas2* mutants, whose mutations provoked a gradual loss of rRNA genes copies (Mozgova et al. 2010). Importantly, lineages 6 and 9 of 20rDNA are the only siblings where the number of rRNA genes remained stable at a low level, whereas all other lineages quickly acquired rRNA genes (Pavlistova et al. 2016). However, our data actually show that rRNA gene copies are increasing progressively in 20rDNA L6F6 (Figure 4A), suggesting that the CNVs are found not only at the level of the TDDOs, but also at the level of the NORs. It remains to be elucidated if a link between the rRNA gene gains and the appearance of TDDO exists and if the same mechanisms are involved. In cancer cells, a loss of rRNA gene copies was associated with genomic instability and hypersensitivity to DNA damage (Wang and Lemos 2017; Xu et al. 2017).

Short-term consequences of gene duplications have been studied in animals, especially in cancer cells, were multiple *de novo* tandem duplication events induce gene CNVs (Wee et al. 2018; Quigley et al. 2018). The 20rDNA L6F6 is an unprecedented opportunity to study the transcriptional behavior of newly duplicated genes. Globally, duplicated genes tend to be more expressed (Figure 2J). In the most recent duplication located on chromosome 2 (TDDO2), 49 out of 97 genes are upregulated more than 2-fold, while 15 out of the 97 can be considered as unexpressed genes. In the short term, the expression of tandem gene duplicates is often greater than two-fold (Loehlin and Carroll 2016). Although we cannot exclude that the detected transcripts come from only one of the duplicated genes, it is more likely that equivalent additive expression occurs for the duplicated genes. During evolution, duplicated gene expression can quickly lead to specialized expression patterns, often in a tissue-specific manner, although a significant number retain correlated transcriptional profiles (Blanc and Wolfe 2004; Guschanski et al. 2017). In our case, we were able to correlate this change in gene expression with the acquisition of increased resistance to different pathogens (Figure 3). Analyzing the expression profile of the genes present in the TDDO2 as well as the ability of the 20rDNA L6F6 to resist to pathogens in future generations will allow us to evaluate if rapid transcriptional regulation will occur.

Plant genomes are rapidly evolving and their capacity to adapt to environmental changes is crucial. Like genome hybridization and TE mobilization, CNV is one important tool of genome evolution (Gabur et al. 2019; Kondrashov 2012; Quadrana et al. 2019). Previous observation and our data demonstrate the importance of systematically detecting CNVs and the presence of chimeric gene formation arising from these duplication events. CNVs can indeed associate with adaptive traits (Kondrashov 2012; Gabur et al. 2019). In our case, we indeed found a potential link between CNV and pathogen resistance (Figure 3 and Suppl. Figure 9). CNVs were already shown to be implicated in nematode resistance in soybean (Cook et al. 2012), but also in potato cultivar genome heterogeneity (Pham et al. 2017). Remarkably, we showed that the CNVs in the 20rDNA line occurred only in a few generations in controlled growing conditions. This last point is particularly interesting in the context of plant breeding. In addition, TDDOs have the potential to create chimeric genes (Figure 5). TDDO events can promote cancer cell formation, *via* the activation of oncogenes (Quigley et al. 2018). In that case, breaking junctions can affect the expression of an oncogene by modifying its regulation by enhancers, for example. In our study, the chimeric genes created are expressed and properly spliced. Although we do not have evidence concerning their potential ability to be translated or if the resultant protein would be stable, it is tempting to speculate that TDDO-mediated chimeric genes can lead to gene novelty as previously described (Chen et al. 2013). Studying the consequences of TDDOs on future generations will certainly shed light on their potential impact on genome evolution and plant adaptation.

## MATERIAL ET METHODS

### Plant materials

Seeds corresponding to the *fas1-4* (SAIL_662_D10) and *fas2-4* (SALK_033228) were previously reported in (Exner et al. 2006). 20rDNA lines depicted in Figure 4C were previously reported in (Pavlistova et al. 2016). For NADs identification, wild-type Col-0 expressing the FIB2:YFP fusion protein was described in (Pontvianne et al. 2013). The 20rDNA L6F5 line was transformed by agroinfiltration to insert a transgene expressing FIB2:YFP fusion protein as described previously (Pontvianne et al. 2013).

### NADs identification

Nuclei and nucleoli were isolated as previously described (Pontvianne et al. 2016a) using a S3 cell sorter (Biorad®). Sorted nuclei or nucleoli were treated with RNase A and proteinase K prior to purification and concentration of the DNA using a phenol/chloroforme purification, followed by two precipitation steps. DNA libraries were generated via the kit Nextera XT DNA sample preparation (Illumina®) according to manufacturer’s instruction, and were then subjected to high throughput paired-end 2X150nt sequencing on a Next-seq 550 apparatus (Illumina ®) at the Bioenvironment sequencing platform (Perpignan University, Perpignan, France). NADs identification was then performed as described in (Carpentier et al. 2018).

### Chromatin-chromatin interactions analyses

Hi-C samples were generated according to (Grob et al. 2014), using *HindIII* as a restriction enzyme. For both wild-type (Columbia-0 accession) and 20rDNA, Hi-C samples were generated in triplicates using 14-day-old seedling populations grown on MS culture media (22°C; 16h light/8h dark). Single ends of paired-end sequencing reads were trimmed to 35 bp using cutadapt (Martin 2011) and aligned separately using bowtie (Langmead et al. 2009) using the following parameters: bowtie -a -v 0 -m 1. After sorting and indexing using samtools (Li et al. 2009), single end alignments were further processed using HiCdat (Schmid et al. 2015). Before mapping read-pairs to genomic bins (10 kb and 50 kb), read pairs were filtered to remove pairs, which paired inwards with less than 1kb and outward with less than 25 kb. The resulting Hi-C matrices were further analyzed in R using HiCdatR package (Schmid et al. 2015) and other in-house R scripts. Hi-C matrices were normalized for sequencing depth (fragment counts per million). To normalize for coverage, we mapped single-end alignments to genomic bins (equal bin size than the corresponding Hi-C matrices) and created bed files indicating the coverage for each genomic bin. Subsequently, the bed files were normalized to obtain read per million values. Hi-C matrices were subsequently converted to coverage-normalized matrices using the R-base scale() function. Principal component analysis (PCA) were performed using f.principle.component.analysis() function from the HiCdatR package using either entire chromosomes or chromosome arms only, excluding pericentromeric regions (Chr1:13Mb-17Mb; Chr2:2Mb-5Mb; Chr3: 11.5Mb-15.5Mb; Chr4: 3Mb-5Mb; Chr5:10Mb;14Mb). Hi-C contact densities were assessed by calculating median interaction frequency of all contacts within 5 times the bin size of a given genomic bin. As input coverage-normalized Hi-C matrices were used. Significant differences between Hi-C data sets were determined by performing individual Student t-tests on each entry of the triplicate Hi-C matrices. Subsequently, we determined 5 categories of changes in contact frequencies (significantly less, less, no change, more, and significantly more). Significant difference matrices were generated using these categories and plotted with an according color code. All plots were generated using R-base.

### Nanopore sequencing and data analyses

Genomic DNA preparation was performed as previously described in (Debladis et al. 2017). After Qubit dosage (dsDNA High Sensitivity (Thermo Fisher Scientific, USA), a second step of DNA purification was performed with the Genomic DNA Clean & Concentrator kit (Zymo Research, USA) and precipitated. A last Qubit dosage was performed before library preparation using the 1D Genomic DNA by ligation kit SQK-LSK109 (Oxford Nanopore Technologies, UK), following manufacturer’s instructions. The R9.5 ONT flow-cell FLO-MIN106D (Oxford Nanopore Technologies, UK) was used.

ONT reads were mapped on the TAIR10 reference genome using minimap2 with -a -Q -map-ont options. The alignment files were converted into bed files using bedtools and the coverage per 100 kb window was calculated using coverageBed. For each 100 kb window the ratio r= 20%rDNA coverage / WT coverage was calculated. The mean m and standard error (SE) were calculated across the entire genome. Differentially covered regions in the 20%rDNA line were defined as regions for which: r>=m+2SE or r<=m-2SE.

### Transcriptome analyses

Total RNA was extracted from four pools of 3-week-old *Arabidopsis* plant leaf tissues of wild-type Col-0 or 20rDNA L6F6 using TRIzol reagent (MRC). Sequencing was performed by the Bioenvironment sequencing platform (Perpignan University, Perpignan, France) using a Nextseq 550 to generate 2 × 75-bp-long reads. Illumina reads from non-stranded, polyA+ RNA deep sequencing libraries were aligned to the *A. thaliana* TAIR10-annotated genome reference using HISAT 2 (Kim et al. 2015). Count the number of reads aligned to each genome coding sequences was performed with HTseq-count (Anders et al. 2015) and differential expression profile analyses with DESeq2 (Bioclite - R package) (Love et al. 2014).

### Methylome analyses

Total genomic DNA was extracted from three pools of 3-week-old *Arabidopsis* plant leaf tissues of wild-type Col-0 or 20rDNA L6F6 using the Illustra Phytopure DNA extraction kit (GE Healthcare®, UK), following manufacturer’s instructions. Bisulfite treatment, libraries preparation and sequencing were performed by the Novogene company (Hong-Kong) using the Illumina® technology. DMRs were identified with Bismark bisulfite mapper (66) and Methylkit (67).

### Immunostaining

Nuclei of *A. thaliana* plants were isolated and immunostained as described (68). An antibody targeting specifically the Phosphorylated γ-H2Ax was used at a dilution of 1:500 in PBS and applied onto a slide pre-coated with nuclei and incubated overnight at 4°C. The nuclei were labeled with anti-rabbit-Alexa Fluor 488 antibodies (Abcam) at a dilution of 1:1000, counterstained with DAPI (4’, 6-diamidino-2-phenylindole), mounted using the Vectashield medium (Vector Laboratories), and analyzed in a Zeiss LSM 700 inverted confocal microscope.

### Bleomycin assay

Sterile were sowed on Petri dishes containing medium ∼ MS (Murashige and Skoog, Duchefa Biochemie M0231) 1X agar: 4.7g / L of MS medium, 5g / L of MES and 10g / L of Plant Agar (Gold Biotechnology) with a pH 5,7. For the treatment, media was supplemented with 1.5 ∼ µg / mL bleomycin to obtain a 10^-6^ M concentration of bleomycin.

### Nematode assay

For the infection assay, ten day-old *A. thaliana* seedlings aseptically grown on modified Knop’s medium at 24°C under 16 h-light/8 h-dark were inoculated with surface-sterilized J2 *H. schachtii* nematodes (69). Four weeks after inoculation, adult females developing on each plant were counted, and the data were analyzed by a modified *t*-test using the Statistical Software Package SAS (P<0.05). Syncytium size measurements was performed 21 days after aseptic inoculation of *A. thaliana* with *H. schachtii* on modified Knop’s in 100 mm petri dishes (70). For each line, 3 pool of 35 single-female syncytia were randomly selected, size was measured and average size for each line determined. Statistically significant differences were determined in a modified *t*-test using the statistical software package SAS (P < 0.05).

### DC3000 infection assay and bacteria counting

Four plants per condition were dip-inoculated using *Pseudomonas* strain DC3000 at 5 x 10^7^ cfu/ml (OD600 of 0.2 corresponds to 10 ^8^cfu/ml) supplemented with 0.02 % Silwet L-77, and immediately placed in chambers with high humidity. Bacterial growth was determined 3 days after infection. For the quantification, infected leaves were harvested, washed for one minute in 70 % (v/v) EtOH and one minute in water. Leaf discs with a diameter of 5 mm have been excised, ground and homogenized in 200 µl of 10 mM MgCl2. Each data point consists of four leaf discs. 10 µl of each homogenate were then plated undiluted and at different dilutions onto NYGA plates. Growth of bacteria was determined after 36 h of incubation at 28°C by colony counting.

### Fluorescence in situ hybridization (FISH), slide preparation, probe labelling

For nuclei isolation for FISH (T5H22, F5I10 ratio) 14 days-old seedlings of WT Col-0 and L6 (generation F6) were used. The seedlings were first fixed in 4% paraformaldehyde for 15 min at RT, chopped with a razor blade in ice cold nucleus isolation buffer (NIB - 0.5 M sucrose; 10 mM EDTA; 2.5 mM DTT; 100 mM KCl; 1 mM spermine; 4 mM spermidine in 10 mM Tris-Cl, pH = 9.5), and filtered through a 50 µm and 30 µm pore-sized disposable filters (CellTrics, Sysmex, Germany). Afterwards, the filtrate was supplemented with 1/10 volume of 10% Triton-X in NIB and centrifuged at 2000 x g for 10 minutes at 4°C. Pellet, containing nuclei, was then resuspended in 1x PBS and nuclei were dripped onto superfrost slides (10 µl / slide). After brief drying at 4°C, cells were fixed in chilled 3:1 methanol / acetic acid mixture for 10 minutes and air dried. After a rinse in 2x SSC, slides were incubated with RNase A (Dnase free, Applichem) in a humid chamber for 45 minutes at 37°C. Slides were then rinsed in 2xSSC and immersed in 10mM HCl for 5 minutes and subsequently in 10mM HCl with 8 µg/ml pepsin for 5 minutes (both at 37°C). After two washes in 2xSSC, slides were dehydrated in ethanol series (70%, 80%, 96%), 2 minutes each, and air-dried.

For pachytene chromosomes, entire inflorescences were fixed in ethanol:glacial acetic acid (3:1) overnight and stored in 70% ethanol at −20°C until use. WT Col-0, L6 (generation F4) and L9 (generation F6) were used. Selected immature flower buds were rinsed in distilled water (2 × 5 min) and in citrate buffer (10 mM sodium citrate, pH 4.8; 2 × 5 min), then anthers were dissected and incubated in an enzyme mix 0.15% each of cellulase (Onozuka R10, Serva), cytohelicase (Sigma), and pectolyase (Duchefa) in citrate buffer (10 mM sodium citrate buffer pH 4.5), at 4°C ON. Individual anthers were put on the microscope slide, disintegrated by the needle in a drop of citrate buffer. Then the suspension was softened by adding 15 to 30 µL of 50% acetic acid and spread by stirring with a needle on a hot plate at 50°C for 1 min. Chromosomes were fixed by adding 100 µL of ethanol:acetic acid (3:1) fixative, the slides were air dried, fixed in 4% formaldehyde dissolved in 2xSSC for 10 min, washed 3×5min in 2xSSC, and treated with RNase and pepsin (5-15min) as described above. After the pepsin treatment, slides were rinsed in 2xSSC, 3×5min and the fixation in paraformaldehyde was repeated, slides washed and dried in the ethanol series, 2 min each, air dried and hybridized with appropriate probes.

Hybridization conditions were used as follows: 50% formamide (Sigma)/10% dextran sulfate/2x SSC solution, denaturation for 3 minutes at 80 °C, overnight incubation at 37 °C. Afterwards, slides were rinsed in 2x SSC at RT, then washed 2×5 min in 50% formamide in 2x SSC and 2×5 min in 2x SSC, (both at 42°C). Slides were blocked in 5% BSA in 4T buffer (4x SSC, 0.05% Tween-20) for 30 minutes at RT. After blocking, incubation with Alexa fluor 594-conjugated streptavidin (#S32356; Life technologies) and Alexa fluor 488-conjugated anti-digoxigenin (#IC75206; R&D systems) was performed for 45 minutes at RT. Antibodies were diluted in blocking buffer (1:200 and 1:10, respectively). Subsequently, slides were washed 2×5 min in 4T buffer and 2×5 min in 2xSSC buffer at 37°C with mild shaking.

Slides were stained with DAPI (4,6-diamidino-2-phenylindole, 2µg/ml) diluted in Vectashield (Vector laboratories), images were acquired using a Zeiss Axioimager Z1 or Olympus BX61, with appropriate filter set (AHF Analysentechnik, http://www.ahf.de/). Images were processed in Photoshop and Image J (https://imagej.nih.gov/ij/) softwares.

The BAC probes (T5H22, F5I10, available at www.arabidopsis.org) were kindly provided by T.Mandáková (CEITEC MU, Brno, CR). Approximately 1 µg of isolated BAC DNA (NucleoBond® Xtra Midi kit, Macherey-Nagel) were labeled by nick translation with biotin-dUTP or digoxigenin-dUTP (Jena Bioscience) using either commercial nick translation labelling kit (#07J00-001; Abbot - labelling was performed according to the manufactureŕs instructions using 5µl of the enzyme mix and 8h labelling time); or by home-made reaction mix (71), containing 5 µL of 10× NT buffer (0.5 M Tris-HCl, pH 7.5, 50 mM MgCl2, and 0.05% BSA), 10mM β-mercaptoethanol, 0.2mM each dATP, dCTP, dGTP and 40 µM dTTP (Jena Bioscience), 0.2 mM labelled-dUTP, 3 µL of DNase I (#10104159001, Roche, diluted to 8 µg/ml) and 5U of DNA polymerase I (#EP0041,Fermentas).The nick translation mixture was incubated at 15°C for ∼ 4h (or longer) to obtain a fragment length of ∼200 to 500 bp. The reaction was stopped by adding 1 µL of 0.5 M EDTA, pH 8.0, and incubation at 65°C for 10 min.

### Quantification of foci in isolated interphase nuclei

In the first analysis we counted the number of hk4 and subNOR4 signals in interphase nuclei and calculated their ratio. Two independent counts were performed and the ratio of hk4 to subNOR4 was evaluated foci in both wild-type cells and L6 cell line. To avoid conscious bias, one of the counts was blind and integrated all the foci into the analysis (singlet, doublet, partially split etc.) The results presented in Suppl. Figure 10 are the average of the two counts. The floral bud nuclei showed were 2, 3 or 4 of hk4 signals. The number and spatial distribution of these signals were evaluated. Total counts are shown in the table in (Suppl. Figure 10B).

## AUTHORS CONTRIBUTION

FP and APC conceived and designed the analysis, APC SG MF TA TM CL EJ FP collected the data, APC SG PZ TB LN MD MM FP contributed data or analysis tools, APC SG NP MM FP performed the analysis, FP wrote the paper, APC SG MM edited the paper, FP acquired funding. TB, VP and TM performed nematode infection assays. LN and TA performed *Pseudomonas DC3000* infection assays. MF and MD performed DNA-FISH analyses. SG performed the HiC analyses.

## AKNOWLEDGEMENTS

The authors are grateful to Rémy Merret and Michèle Laudié for Illumina sequencing and lab members for fruitful discussions. We also thank the flow cytometry facility, the microscopic facility and the sequencing facility of Perpignan University *Via Domitia* Bioenvironnement (Perpignan, France).

## DECLARATION

The author(s) declare(s) that they have no competing interests”.

## Figure Legends

**Supplemental Figure 1:**

**A-B**. Chromosome plots displaying the relative enrichment of a given genomic segment with the nucleolus in WT Col-0 (A) and in the 20rDNA L6F6 (B). The y-axis displays the fold change between the nuclear *versus* the nucleolar DNA. Each dot represents a 100kb window. Nucleolus-enriched genomic regions above the threshold (red-dotted line) are colored in red.

**Supplemental Figure 2:**

**A-B.** HiC matrix displaying the chromatin-chromatin interactions identified by the chromosome capture conformation technique in wild-type Col-0 (A) and the 20rDNA L6F6 (B). Enriched interactions are displayed in yellow.

**Supplemental Figure 3:**

Eigenvector (in green) of the Col-0 wild-type Hi-C data set and annotation of the TDDO. TEs density is marked in red and the violet square show the TDDOs location.

**Supplemental Figure 4:**

**A.** Representative confocal images showing the immunolocalization of the P-γH2Ax foci (green) in WT Col-0 (top panels) or in 20rDNA F6L6 (bottom panels) isolated nuclei. The Blue staining correspond to the DNA labeled with DAPI. **B.** WT Col-0 and 20rDNA L6F6 seeds were grown for 18 days on an MS plate with or without Bleomycine at 10^-5^M. Individual plant growth was then measured using their fresh weights.

**Supplemental Figure 5:**

**A.** Representation of the chromosomal position of genes and TEs of differentially accumulated transcripts in the 20rDNA L6F6 *versus* WT Col-0 using *Circos* (72). Up-regulated TEs are in green, Up-regulated genes are in red and down-regulated genes are in blue. **B**. QPCR analyses of differentially accumulated genes. Their relative accumulation was analyzed against the *ACTIN7* (red bars) or the *HC7* (blue bars) household genes.

**Supplemental Figure 6:**

**A.** Venn diagram showing the relation between genes that lost their association with the nucleolus in the 20rDNA L6F6 (lost NAD-genes in 20rDNA), Genes that are upregulated in 20rDNA L6F6 with an adjusted *p-*value < 0.01 and a log2FoldChange > 2 (Genes UP in 20rDNA) and with hypomethylated genes in 20rDNA L6F6. **B**. Venn diagram showing the relation between genes that gain association with the nucleolus in the 20rDNA L6F6 (Acquired NAD-genes in 20rDNA), Genes that are down-regulated in 20rDNA L6F6 with an adjusted *p-*value < 0.01 and a log2FoldChange > 2 (Genes Down in 20rDNA) and with hypermethylated genes in 20rDNA L6F6. **C**. Venn diagram showing the relation between TEs that lost their association with the nucleolus in the 20rDNA L6F6 (lost NAD-TEs in 20rDNA), TEs that are upregulated in 20rDNA L6F6 with an adjusted *p-*value < 0.01 and a log2FoldChange > 2 (TEs UP in 20rDNA) and with hypomethylated TEs in 20rDNA L6F6. **D-E.** Venn diagrams representing the proportion of duplicated genes (D) or TEs (E) that accumulated a higher level of transcripts, using an adjusted *p* value < 0.01 and a log2FoldChange > 1.5

**Supplemental Figure 7:**

**A-F**. Dot plots showing the relative methylation levels in CG, CHG and CHH contexts in WT Col-0 or in 20rDNA L6F6. The analysis was performed by genome-wide bisulfite sequencing. 2 WT Col-0 and 2 20rDNA L6F6 (labelled 20) replicates (1 and 2) are shown. The average methylation between both replicates is shown (labelled ave.). The analysis was performed with All genes (A-B), with Upregulated genes with an adjusted *p-*value < 0.01 and a log2FoldChange > 1.5 (C-D) and with the Duplicated genes (E-F). For this analysis, we separated Gene body sequences (top panels A, C and E) from the flanking sequences of the genes (5’ and 3’ – bottom panels B, D and F). *(p<0,00025); **(p<5.10^-10^); ***(p<2.10^-16^) calculated using a Wilcoxon test.

**Supplemental Figure 8:**

**A-D**. Dot plots showing the relative methylation levels in CG, CHG and CHH contexts in WT Col-0 or in 20rDNA L6F6. The analysis was performed by genome-wide bisulfite sequencing. 2 WT Col-0 and 2 20rDNA L6F6 (labelled 20) replicates (1 and 2) are shown. The average methylation between both replicates is shown (labelled ave.). The analysis was performed with All TEs (A-B) and with the Duplicated TEs (C-D). For this analysis, we separated TE body sequences (top panels A and C) from the flanking sequences of the TEs (5’ and 3’ – bottom panels B and D). *(p<0,00025); **(p<5.10^-10^); ***(p<2.10^-16^) calculated using a Wilcoxon test.

**Supplemental Figure 9:**

**A-B**. Genome browser (IGV - (73)) screenshot showing examples of Up-genes in 20rDNA L6F6 in duplicated regions. DNA coverage was obtained by nanopore sequencing for WT Col-0 (green) and for 20rDNA L6F6 (blue). Reads from RNA sequencing are shown for WT Col-0 (red) and for 20rDNA L6F6 (yellow). On the bottom, genes location is represented in blue and TEs in green. Gene names are surrounded in blue (non-significant) and in red (significant) when they are differentially accumulated transcripts with an adjusted *p-*value < 0.01 and a log2FoldChange > 1.5. This value is noted in red. Two locations are presented: the left border of the TDDO3 (A) and portion of the TDDO4 (B).

**Supplemental Figure 10:**

**A-B.** DNA-FISH analyses of a portion of the TDDO4 in leaf nuclei. **A.** Two *loci* present on kr4s, distant by 1.6 Mb: one present on the TDDO4 (BAC T5H22 – green) and one absent of the duplicated region (BAC F5I10 – red). Three leaf nuclei from WT Col-0 and 20rDNA Line 6 are shown and the ratio of green versus red signals as well as its ratio noted in the figure. **B.** In that case, the TDDO4 locus (BAC T5H22 – green) was analyzed in floral bud nuclei and four representatives images are presented in the figure. DNA is labelled with DAPI stain (Top panel, with DAPI in grey). Between 58 and 77 nuclei per genotype (WT Col-0, rDNA20 L6 and L9) and the distribution of each nuclei found per genotype is presented in the table.

**Supplemental Figure 11:**

Distribution of the TDDOs along the *A. thaliana* chromosomes and analyses of the breaking junctions. Genes present in the breaking junction and their orientation are represented by an arrow in green (for the 5’ of the TDDO) and in red (for the 3’ of the TDDO).

**Supplemental Figure 12:**

A-D. PCR performed with primers flanking the breaking junctions of the TDDO1 (A), TDDO3 (B) and TDDO4 (C) in the wild-type Col-0, the two mutants *fas1-4* and *fas2-4*, the 20rDNA lines L6 (generations F6 and F8) and L9 (generation F7). For the TDDO1, the cDNA amplified for shorter using cDNA versus gDNA template due to the correct splicing of the three introns, as confirmed by Sanger sequencing The locus encoding the elongation factor *EF1α* was used as a loading control in (D).

## REFERENCES

1. G. P. Copenhaver, C. S. Pikaard, RFLP and physical mapping with an rDNA-specific endonuclease reveals that nucleolus organizer regions of Arabidopsis thaliana adjoin the telomeres on chromosomes 2 and 4. *Plant J*. Cell Mol. Biol. 9, 259–272 (1996).

2. I. Grummt, G. Langst, Epigenetic control of RNA polymerase I transcription in mammalian cells. Biochim. Biophys. Acta 1829, 393–404 (2013).

3. F. Pontvianne, et al., Subnuclear partitioning of rRNA genes between the nucleolus and nucleoplasm reflects alternative epiallelic states. Genes Dev. 27, 1545–1550 (2013).

4. F. Pontvianne, et al., Histone methyltransferases regulating rRNA gene dose and dosage control in Arabidopsis. Genes Dev. 26, 945–957 (2012).

5. F. Pontvianne, et al., Nucleolin is required for DNA methylation state and the expression of rRNA gene variants in Arabidopsis thaliana. PLoS Genet. 6, e1001225 (2010).

6. T. Kobayashi, Regulation of ribosomal RNA gene copy number and its role in modulating genome integrity and evolutionary adaptability in yeast. Cell. Mol. Life Sci. CMLS 68, 1395–1403 (2011).

7. E. B. Dopman, D. L. Hartl, A portrait of copy-number polymorphism in Drosophila melanogaster. Proc. Natl. Acad. Sci. U. S. A. 104, 19920–19925 (2007).

8. J. G. Gibbons, A. T. Branco, S. A. Godinho, S. Yu, B. Lemos, Concerted copy number variation balances ribosomal DNA dosage in human and mouse genomes. Proc. Natl. Acad. Sci. U. S. A. 112, 2485–2490 (2015).

9. F. A. Rabanal, et al., Unstable Inheritance of 45S rRNA Genes in Arabidopsis thaliana. G3 Bethesda Md 7, 1201–1209 (2017).

10. Q. Long, et al., Massive genomic variation and strong selection in Arabidopsis thaliana lines from Sweden. Nat. Genet. 45, 884–890 (2013).

11. S. Paredes, A. T. Branco, D. L. Hartl, K. A. Maggert, B. Lemos, Ribosomal DNA deletions modulate genome-wide gene expression: “rDNA-sensitive” genes and natural variation. PLoS Genet. 7, e1001376 (2011).

12. B. Lemos, L. O. Araripe, P. Fontanillas, D. L. Hartl, Dominance and the evolutionary accumulation of cis- and trans-effects on gene expression. Proc. Natl. Acad. Sci. U. S. A. 105, 14471–14476 (2008).

13. E. Ramirez-Parra, C. Gutierrez, The many faces of chromatin assembly factor 1. Trends Plant Sci. 12, 570–576 (2007).

14. I. Mozgova, P. Mokros, J. Fajkus, Dysfunction of chromatin assembly factor 1 induces shortening of telomeres and loss of 45S rDNA in Arabidopsis thaliana. Plant Cell 22, 2768–2780 (2010).

15. V. Pavlistova, et al., Phenotypic reversion in fas mutants of Arabidopsis thaliana by reintroduction of FAS genes: variable recovery of telomeres with major spatial rearrangements and transcriptional reprogramming of 45S rDNA genes. Plant J. Cell Mol. Biol. 88, 411–424 (2016).

16. C. Bersaglieri, R. Santoro, Genome Organization in and around the Nucleolus. Cells 8 (2019).

17. A. Nemeth, et al., Initial genomics of the human nucleolus. PLoS Genet. 6, e1000889 (2010).

18. S. van Koningsbruggen, et al., High-resolution whole-genome sequencing reveals that specific chromatin domains from most human chromosomes associate with nucleoli. Mol. Biol. Cell 21, 3735–3748 (2010).

19. F. Pontvianne, et al., Identification of Nucleolus-Associated Chromatin Domains Reveals a Role for the Nucleolus in 3D Organization of the A. thaliana Genome. Cell Rep. 16, 1574–1587 (2016).

20. A. Picart-Picolo, N. Picault, F. Pontvianne, Ribosomal RNA genes shape chromatin domains associating with the nucleolus. Nucl. Austin Tex, 1–6 (2019).

21. S. A. Quinodoz, et al., Higher-Order Inter-chromosomal Hubs Shape 3D Genome Organization in the Nucleus. Cell 174, 744–757.e24 (2018).

22. F. Pontvianne, M. Boyer-Clavel, J. Saez-Vasquez, Fluorescence-Activated Nucleolus Sorting in Arabidopsis. Methods Mol. Biol. Clifton NJ 1455, 203–211 (2016).

23. M.-C. Carpentier, A. Picart-Picolo, F. Pontvianne, A Method to Identify Nucleolus-Associated Chromatin Domains (NADs). Methods Mol. Biol. Clifton NJ 1675, 99–109 (2018).

24. C. Chandrasekhara, G. Mohannath, T. Blevins, F. Pontvianne, C. S. Pikaard, Chromosome-specific NOR inactivation explains selective rRNA gene silencing and dosage control in Arabidopsis. Genes Dev. 30, 177–190 (2016).

25. A. Himmelbach, et al., Discovery of multi-megabase polymorphic inversions by chromosome conformation capture sequencing in large-genome plant species. *Plant J*. Cell Mol. Biol. 96, 1309–1316 (2018).

26. S. Grob, M. W. Schmid, U. Grossniklaus, Hi-C analysis in Arabidopsis identifies the KNOT, a structure with similarities to the flamenco locus of Drosophila. Mol. Cell 55, 678– 693 (2014).

27. E. Lieberman-Aiden, et al., Comprehensive mapping of long-range interactions reveals folding principles of the human genome. Science 326, 289–293 (2009).

28. L. Zapata, et al., Chromosome-level assembly of Arabidopsis thaliana Ler reveals the extent of translocation and inversion polymorphisms. Proc. Natl. Acad. Sci. U. S. A. 113, E4052–4060 (2016).

29. C. Charbonnel, M. E. Gallego, C. I. White, Xrcc1-dependent and Ku-dependent DNA double-strand break repair kinetics in Arabidopsis plants. *Plant J*. Cell Mol. Biol. 64, 280–290 (2010).

30. M. J. E. Wubben, J. Jin, T. J. Baum, Cyst nematode parasitism of Arabidopsis thaliana is inhibited by salicylic acid (SA) and elicits uncoupled SA-independent pathogenesis-related gene expression in roots. Mol. Plant-Microbe Interact. MPMI 21, 424–432 (2008).

31. J.-Y. Yang, et al., betaC1, the pathogenicity factor of TYLCCNV, interacts with AS1 to alter leaf development and suppress selective jasmonic acid responses. Genes Dev. 22, 2564–2577 (2008).

32. P. L. Nurmberg, et al., The developmental selector AS1 is an evolutionarily conserved regulator of the plant immune response. Proc. Natl. Acad. Sci. U. S. A. 104, 18795–18800 (2007).

33. Y.-H. Yeh, Y.-H. Chang, P.-Y. Huang, J.-B. Huang, L. Zimmerli, Enhanced Arabidopsis pattern-triggered immunity by overexpression of cysteine-rich receptor-like kinases. Front. Plant Sci. 6, 322 (2015).

34. S. Newman, K. E. Hermetz, B. Weckselblatt, M. K. Rudd, Next-generation sequencing of duplication CNVs reveals that most are tandem and some create fusion genes at breakpoints. Am. J. Hum. Genet. 96, 208–220 (2015).

35. K. V. Krasileva, The role of transposable elements and DNA damage repair mechanisms in gene duplications and gene fusions in plant genomes. Curr. Opin. Plant Biol. 48, 18–25 (2019).

36. G. Blanc, A. Barakat, R. Guyot, R. Cooke, M. Delseny, Extensive duplication and reshuffling in the Arabidopsis genome. Plant Cell 12, 1093–1101 (2000).

37. Y. Henry, M. Bedhomme, G. Blanc, History, protohistory and prehistory of the Arabidopsis thaliana chromosome complement. Trends Plant Sci. 11, 267–273 (2006).

38. W.-B. Jiao, K. Schneeberger, Chromosome-level assemblies of multiple *Arabidopsis thaliana* accessions reveal hotspots of genomic rearrangements. bioRxiv, 738880 (2019).

39. A. Fulgione, A. M. Hancock, Archaic lineages broaden our view on the history of Arabidopsis thaliana. New Phytol. 219, 1194–1198 (2018).

40. J. Zhang, T. Zuo, T. Peterson, Generation of tandem direct duplications by reversed-ends transposition of maize ac elements. PLoS Genet. 9, e1003691 (2013).

41. J. O. Nelson, G. J. Watase, N. Warsinger-Pepe, Y. M. Yamashita, Mechanisms of rDNA Copy Number Maintenance. Trends Genet. TIG 35, 734–742 (2019).

42. K. D. Tartof, Unequal mitotic sister chromatid exchange and disproportionate replication as mechanisms regulating ribosomal RNA gene redundancy. Cold Spring Harb. Symp. Quant. Biol. 38, 491–500 (1974).

43. K. D. Tartof, Unequal mitotic sister chromatin exchange as the mechanism of ribosomal RNA gene magnification. Proc. Natl. Acad. Sci. U. S. A. 71, 1272–1276 (1974).

44. M. Wang, B. Lemos, Ribosomal DNA copy number amplification and loss in human cancers is linked to tumor genetic context, nucleolus activity, and proliferation. PLoS Genet. 13, e1006994 (2017).

45. B. Xu, et al., Ribosomal DNA copy number loss and sequence variation in cancer. PLoS Genet. 13, e1006771 (2017).

46. Y. Wee, T. Wang, Y. Liu, X. Li, M. Zhao, A pan-cancer study of copy number gain and up-regulation in human oncogenes. Life Sci. 211, 206–214 (2018).

47. D. A. Quigley, et al., Genomic Hallmarks and Structural Variation in Metastatic Prostate Cancer. Cell 174, 758–769.e9 (2018).

48. D. W. Loehlin, S. B. Carroll, Expression of tandem gene duplicates is often greater than twofold. Proc. Natl. Acad. Sci. U. S. A. 113, 5988–5992 (2016).

49. G. Blanc, K. H. Wolfe, Functional divergence of duplicated genes formed by polyploidy during Arabidopsis evolution. Plant Cell 16, 1679–1691 (2004).

50. K. Guschanski, M. Warnefors, H. Kaessmann, The evolution of duplicate gene expression in mammalian organs. Genome Res. 27, 1461–1474 (2017).

51. I. Gabur, H. S. Chawla, R. J. Snowdon, I. A. P. Parkin, Connecting genome structural variation with complex traits in crop plants. TAG Theor. Appl. Genet. Theor. Angew. Genet. 132, 733–750 (2019).

52. F. A. Kondrashov, Gene duplication as a mechanism of genomic adaptation to a changing environment. Proc. Biol. Sci. 279, 5048–5057 (2012).

53. L. Quadrana, et al., Transposition favors the generation of large effect mutations that may facilitate rapid adaption. Nat. Commun. 10, 3421 (2019).

54. D. E. Cook, et al., Copy number variation of multiple genes at Rhg1 mediates nematode resistance in soybean. Science 338, 1206–1209 (2012).

55. G. M. Pham, et al., Extensive genome heterogeneity leads to preferential allele expression and copy number-dependent expression in cultivated potato. *Plant J*. Cell Mol. Biol. 92, 624–637 (2017).

56. S. Chen, B. H. Krinsky, M. Long, New genes as drivers of phenotypic evolution. Nat. Rev. Genet. 14, 645–660 (2013).

57. V. Exner, P. Taranto, N. Schonrock, W. Gruissem, L. Hennig, Chromatin assembly factor CAF-1 is required for cellular differentiation during plant development. Dev. Camb. Engl. 133, 4163–4172 (2006).

58. M. Martin, Cutadapt removes adapter sequences from high-throughput sequencing reads. EMBnet.journal 17, 3 (2011).

59. B. Langmead, C. Trapnell, M. Pop, S. L. Salzberg, Ultrafast and memory-efficient alignment of short DNA sequences to the human genome. Genome Biol. 10, R25 (2009).

60. H. Li, et al., The Sequence Alignment/Map format and SAMtools. Bioinforma. Oxf. Engl. 25, 2078–2079 (2009).

61. M. W. Schmid, S. Grob, U. Grossniklaus, HiCdat: a fast and easy-to-use Hi-C data analysis tool. BMC Bioinformatics 16, 277 (2015).

62. E. Debladis, C. Llauro, M.-C. Carpentier, M. Mirouze, O. Panaud, Detection of active transposable elements in Arabidopsis thaliana using Oxford Nanopore Sequencing technology. BMC Genomics 18, 537 (2017).

63. D. Kim, B. Langmead, S. L. Salzberg, HISAT: a fast spliced aligner with low memory requirements. Nat. Methods 12, 357–360 (2015).

64. S. Anders, P. T. Pyl, W. Huber, HTSeq--a Python framework to work with high-throughput sequencing data. Bioinforma. Oxf. Engl. 31, 166–169 (2015).

65. M. I. Love, W. Huber, S. Anders, Moderated estimation of fold change and dispersion for RNA-seq data with DESeq2. Genome Biol. 15, 550 (2014).

66. F. Krueger, S. R. Andrews, Bismark: a flexible aligner and methylation caller for Bisulfite-Seq applications. Bioinforma. Oxf. Engl. 27, 1571–1572 (2011).

67. A. Akalin, et al., methylKit: a comprehensive R package for the analysis of genome-wide DNA methylation profiles. Genome Biol. 13, R87 (2012).

68. N. Durut, et al., A duplicated NUCLEOLIN gene with antagonistic activity is required for chromatin organization of silent 45S rDNA in Arabidopsis. Plant Cell 26, 1330–1344 (2014).

69. T. J. Baum, M. J. Wubben, K. A. Hardyy, H. Su, S. R. Rodermel, A Screen for Arabidopsis thaliana Mutants with Altered Susceptibility to Heterodera schachtii. J. Nematol. 32, 166–173 (2000).

70. T. Hewezi, et al., Cellulose binding protein from the parasitic nematode Heterodera schachtii interacts with Arabidopsis pectin methylesterase: cooperative cell wall modification during parasitism. Plant Cell 20, 3080–3093 (2008).

71. T. Mandakova, M. A. Lysak, Chromosomal phylogeny and karyotype evolution in x=7 crucifer species (Brassicaceae). Plant Cell 20, 2559–2570 (2008).

