## Supplementary material for "Low ribosomal RNA genes copy number provoke genomic instability and chromosomal segment duplication events that modify global gene expression and plant-pathogen response": Suppl. Figures

Suppl. Figure 1

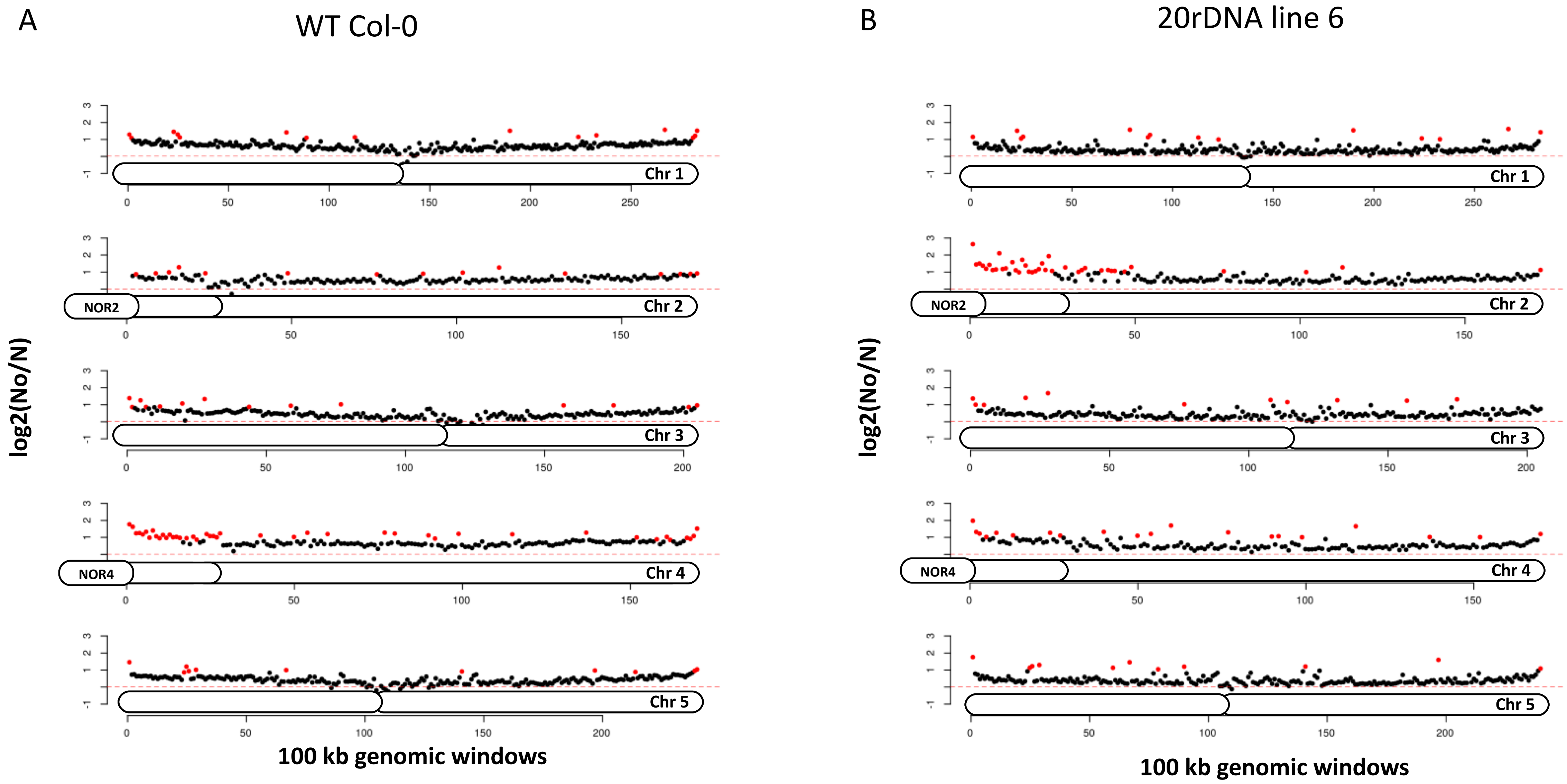

Suppl. Figure 2

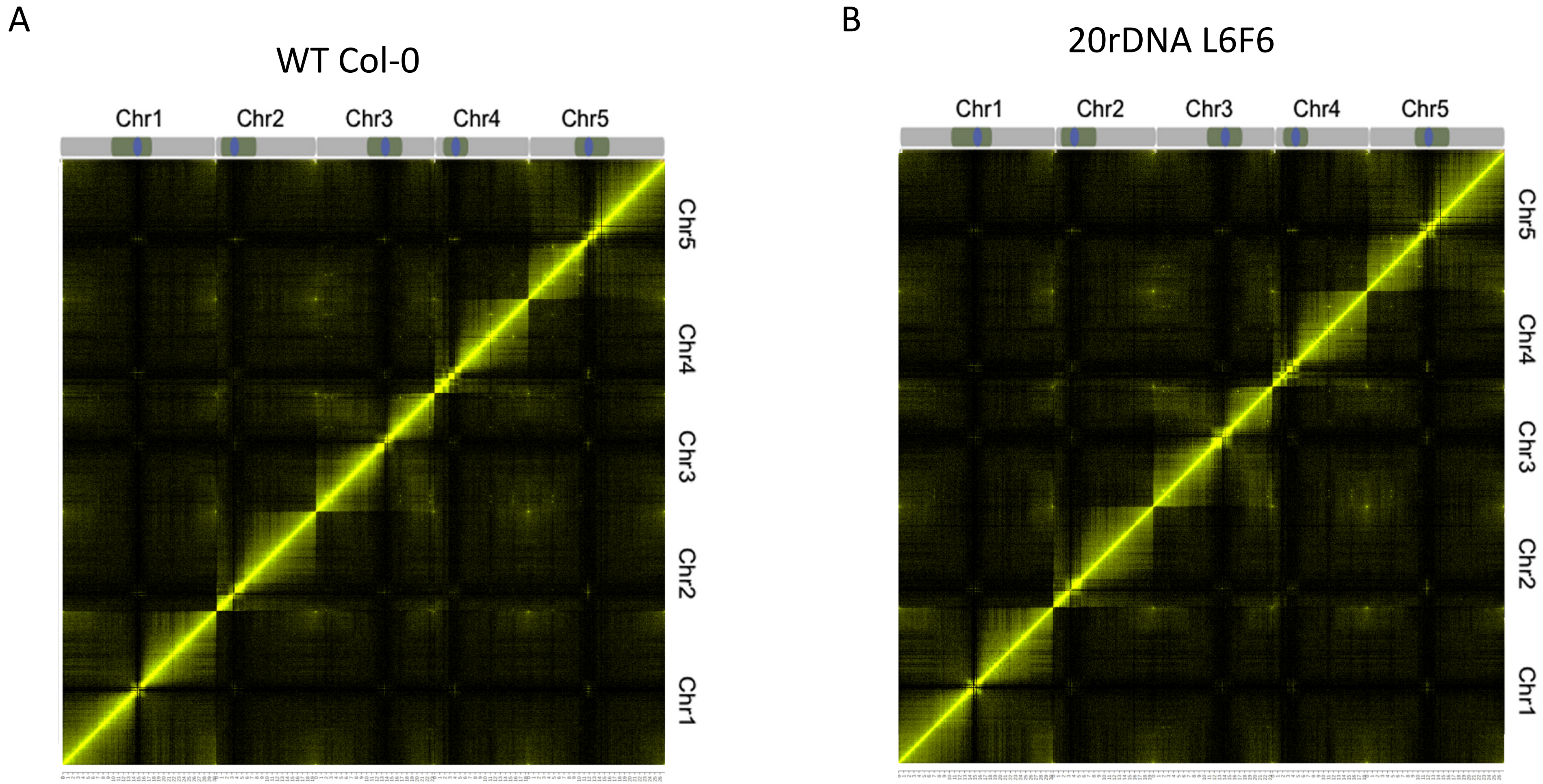

### Suppl. Figure 3

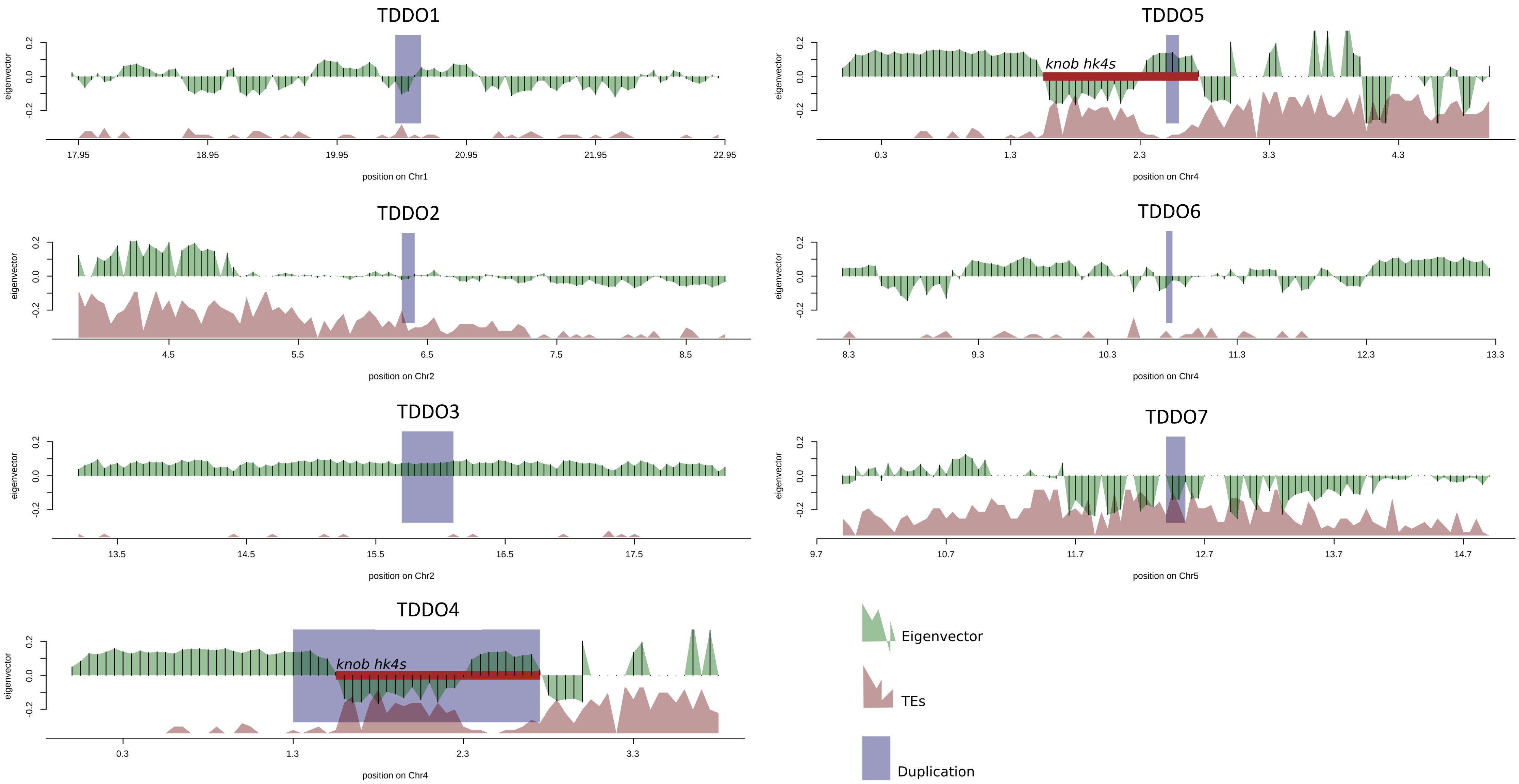

Suppl. Figure 4

A

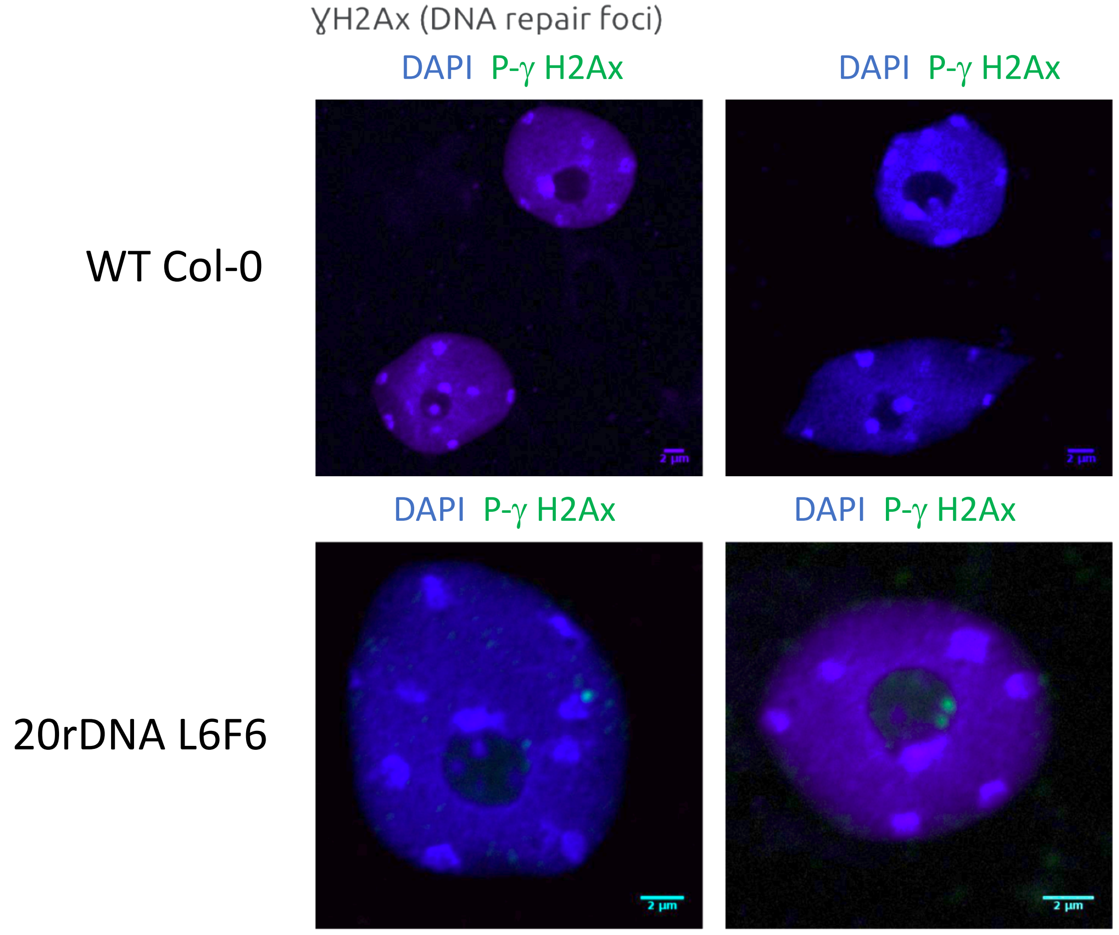


B

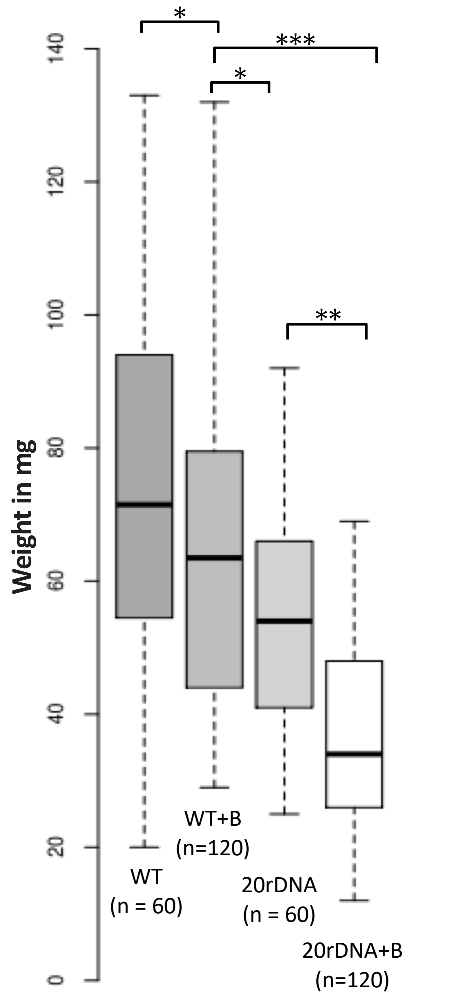


**Paired Student's t test**
(two sided, non equal variances)

* = p value < 0.001
** = p value < 0.00001 (e-5)
*** = p value < 0.0000001 (e-7)

Suppl. Figure 5

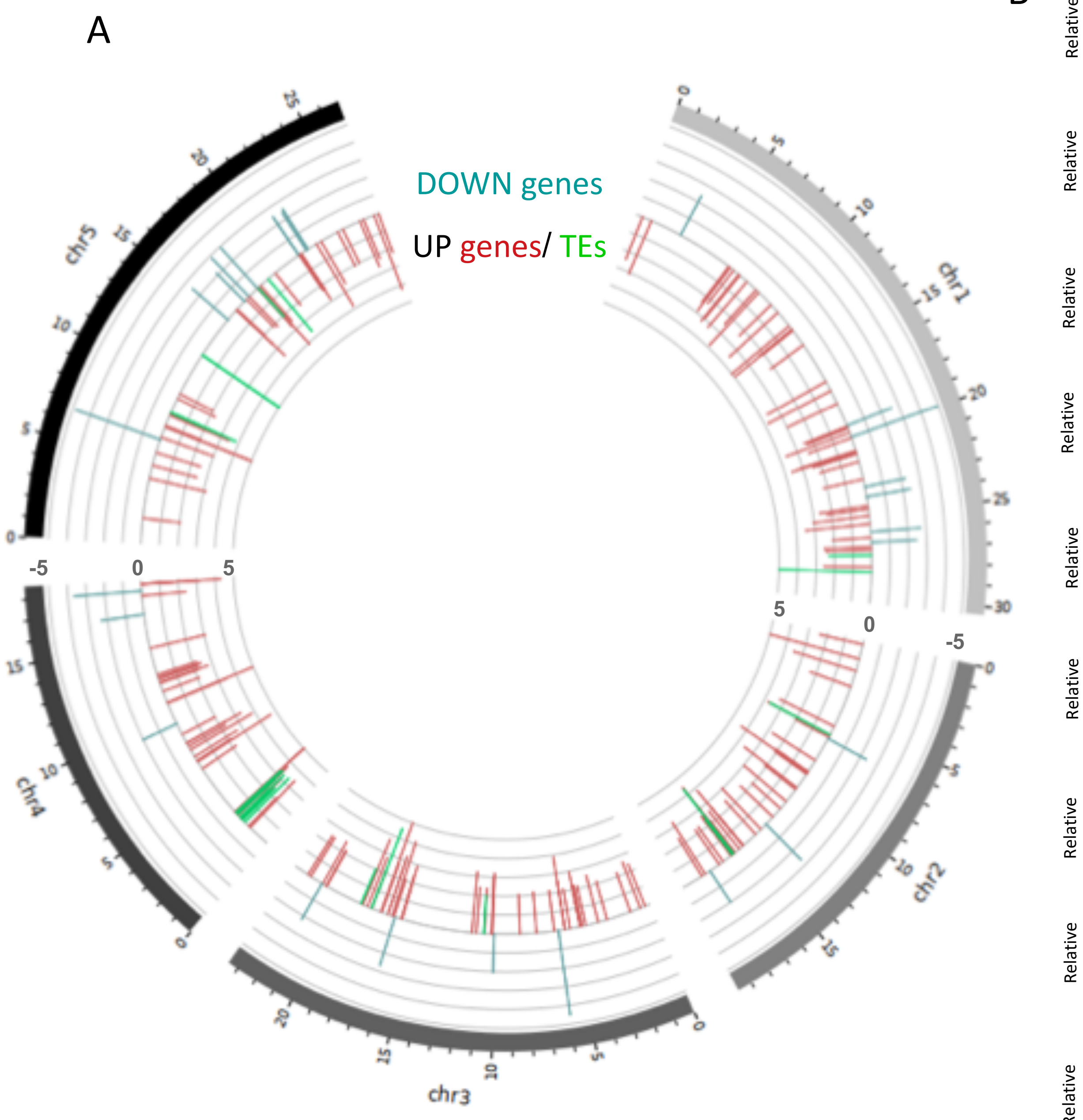


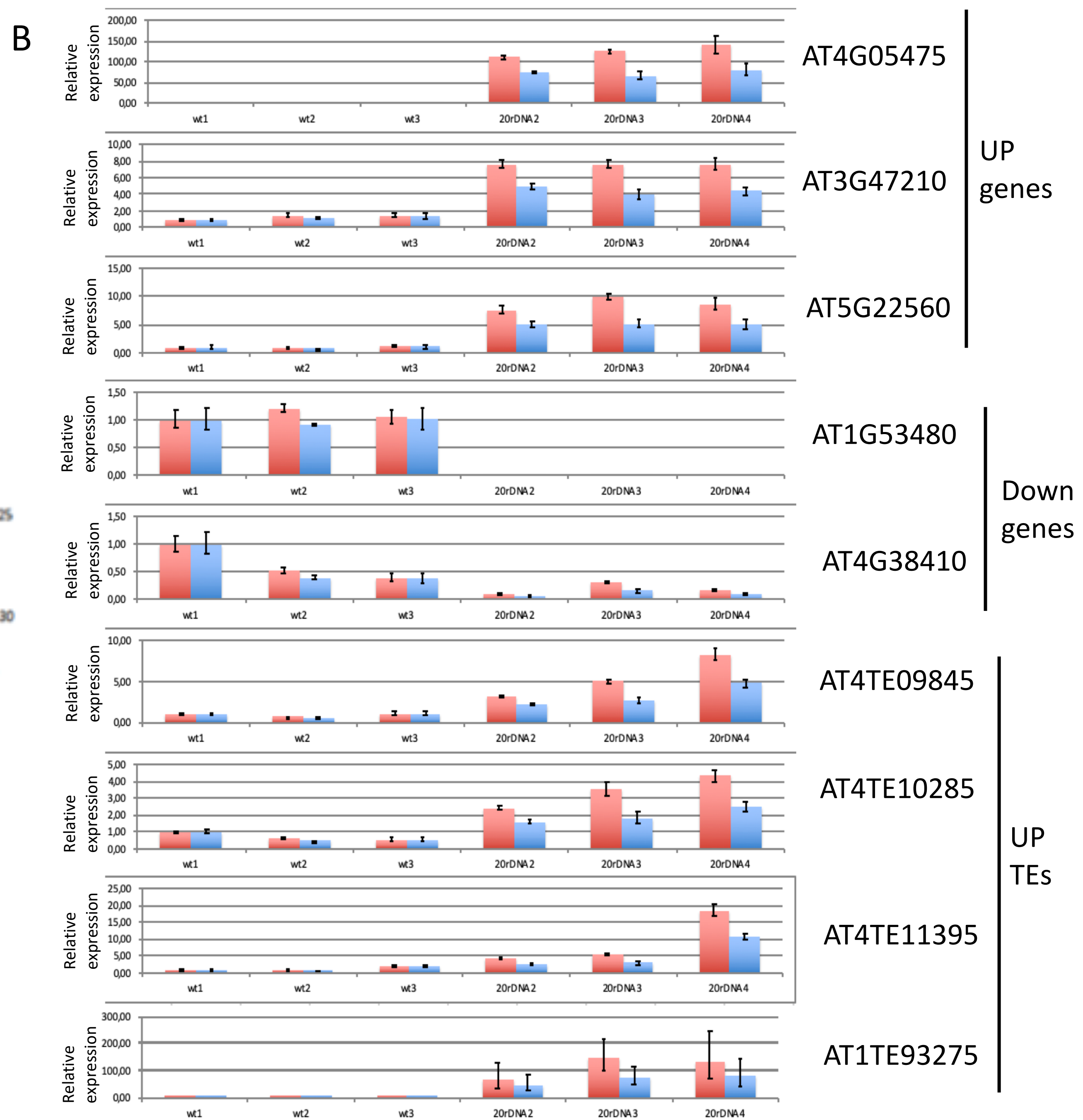

Suppl. Figure 6

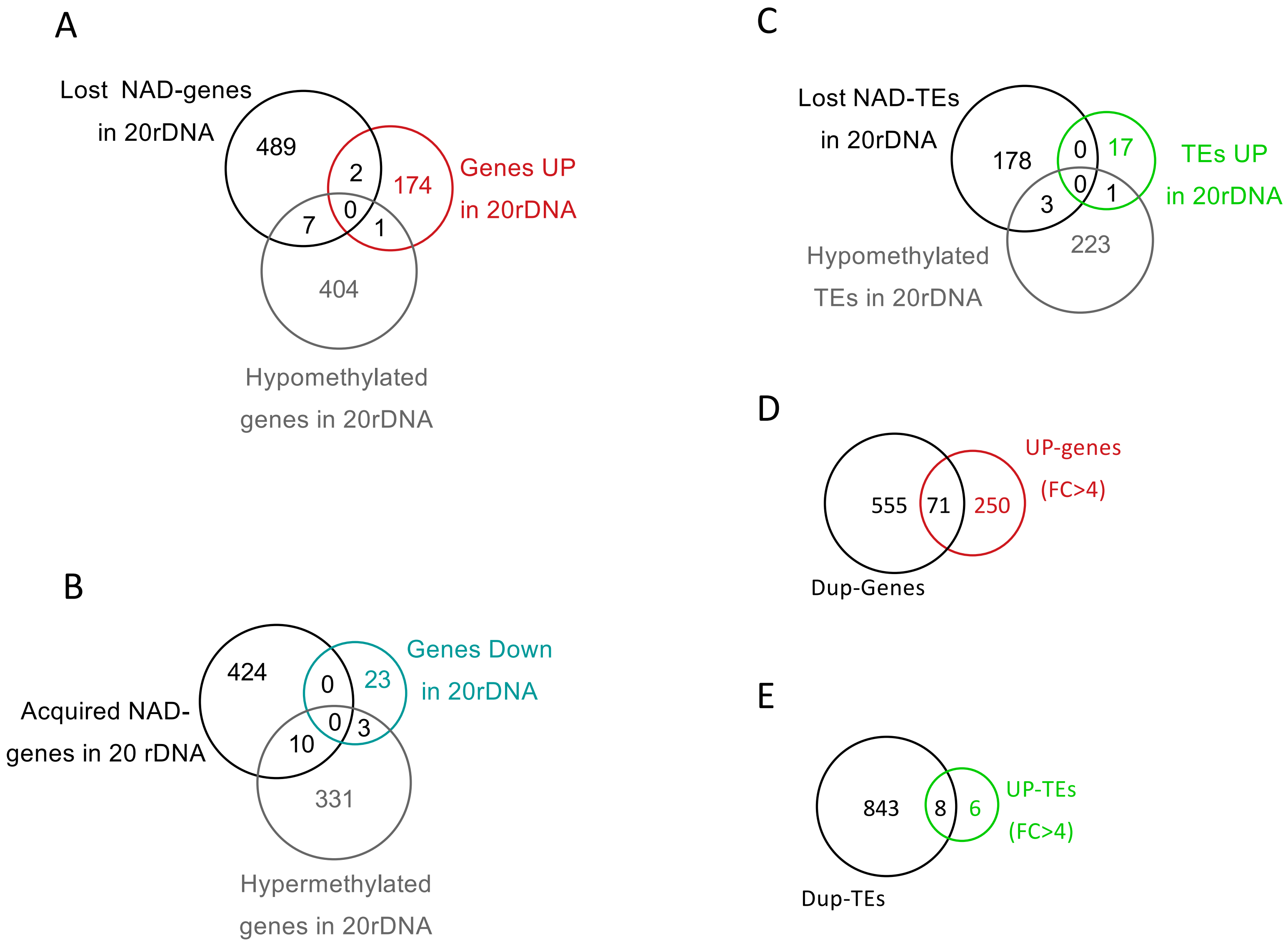

Suppl. Figure 7

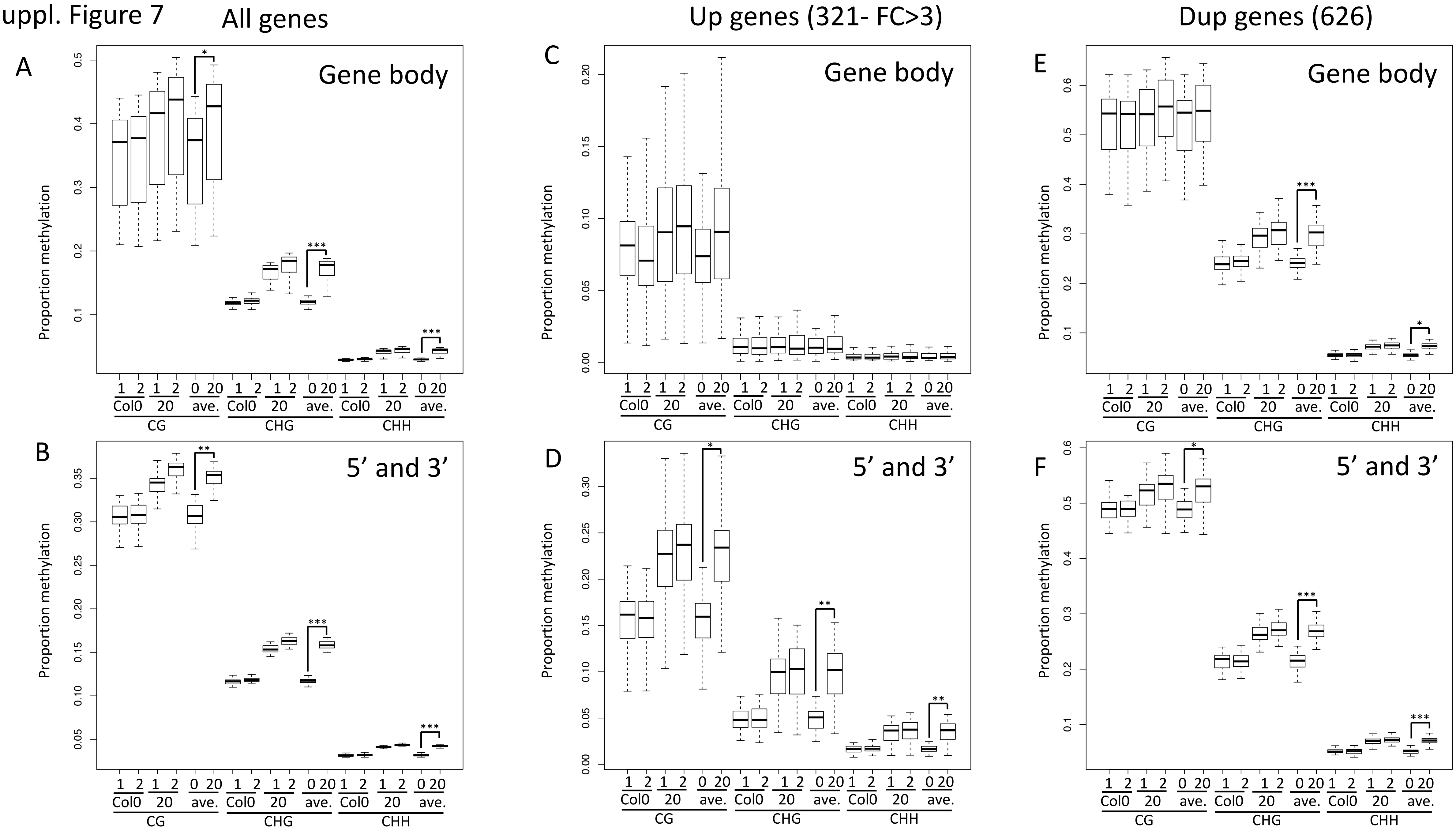

Suppl. Figure 8

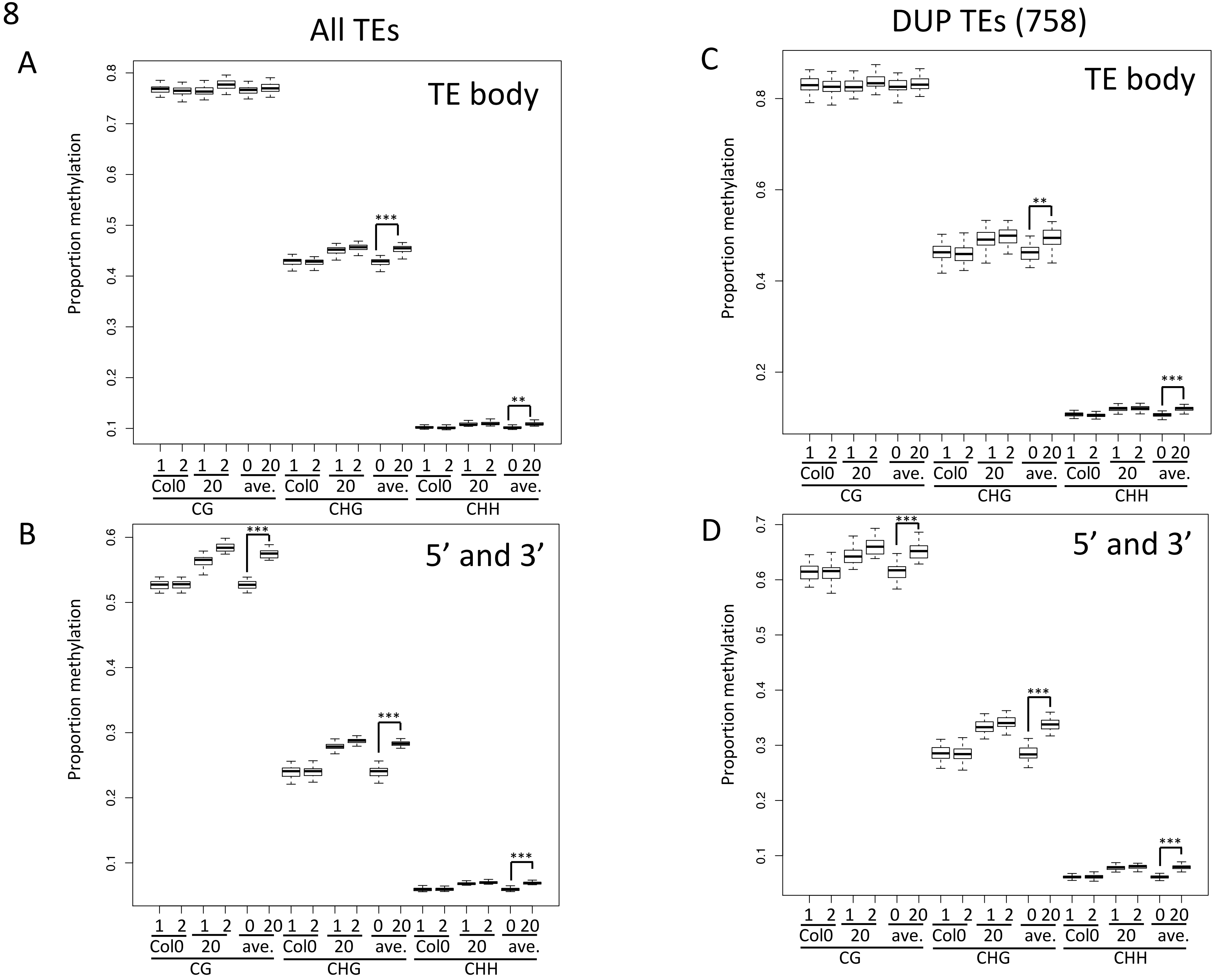

Suppl. Figure 9

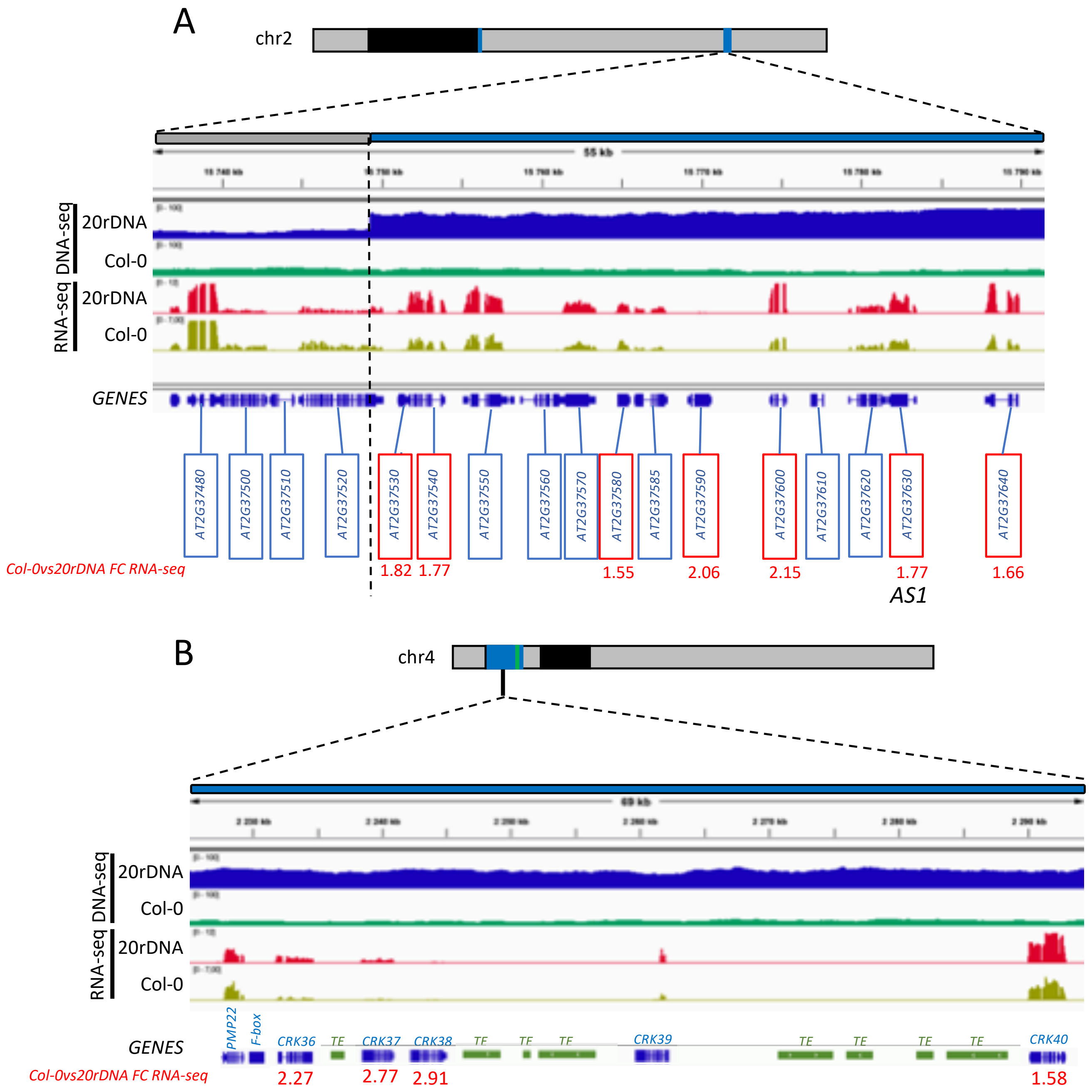

Suppl. Figure 10

A

**LEAF NUCLEI**

WT

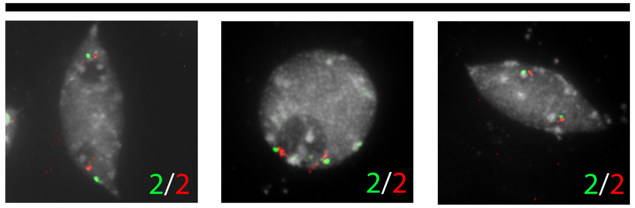


hk to subNOR foci number ratio - 1.06 (n = 30)

L6

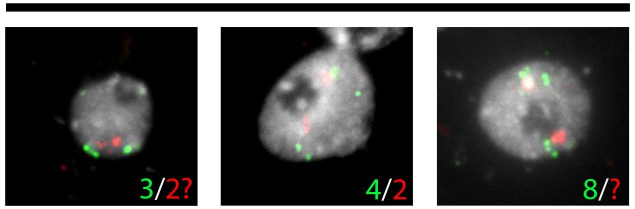


hk to subNOR foci number ratio - 1.47 (n = 42)

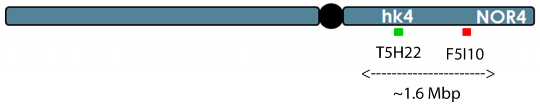


B

**FLORAL BUD NUCLEI**

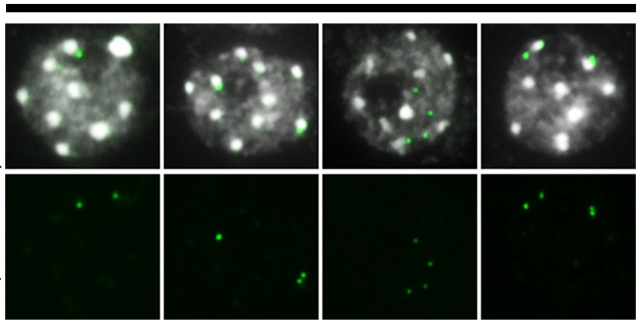


TYPE 1 TYPE 2 TYPE 3

| | TYPE 1 | TYPE 2 | TYPE 3 | total |
|---|---|---|---|---|
| WT | 77 | 0 | 0 | 77 |
| L6 | 8 | 24 | 34 | 66 |
| L9 | 56 | 2 | 0 | 58 |

* in WT also some foci appear large, as if there were double dots, but not separated as in L6

Suppl. Figure 11

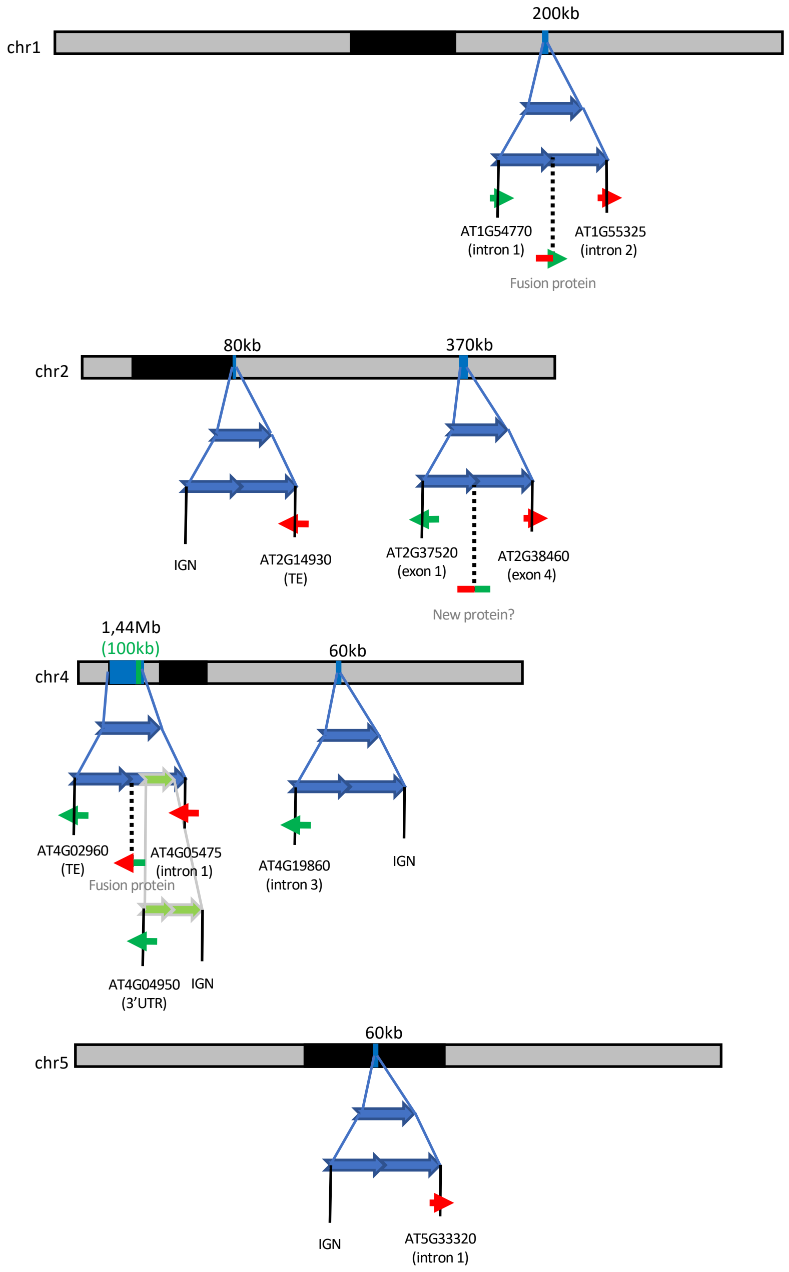

Suppl. Figure 12

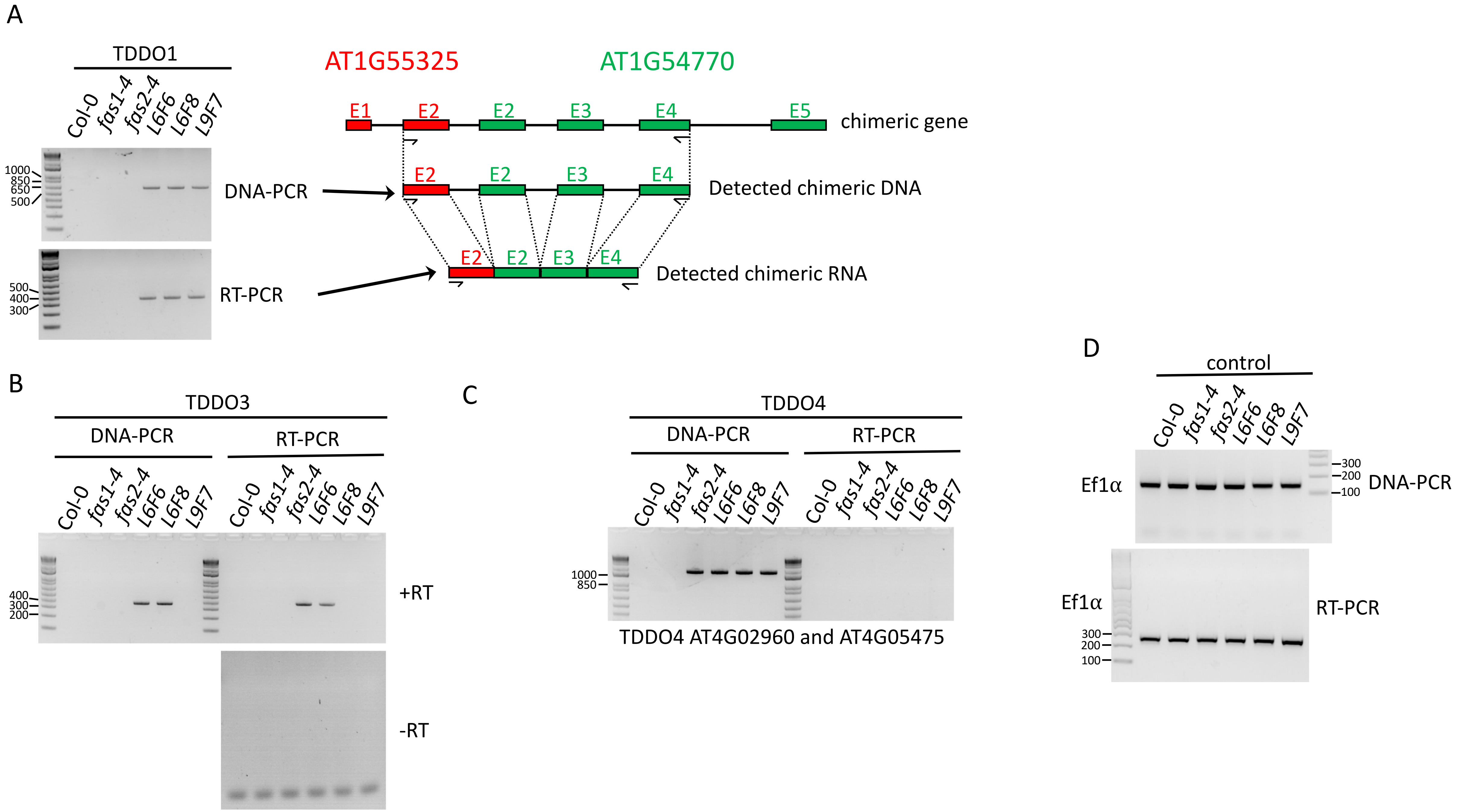
